# Constitutive activation of cellular immunity underlies the evolution of resistance to infection

**DOI:** 10.1101/2020.05.01.072413

**Authors:** Alexandre B. Leitão, Ramesh Arunkumar, Jonathan P. Day, Emma M. Geldman, Francis M. Jiggins

## Abstract

Organisms rely on inducible and constitutive immune defences to combat infection. Constitutive immunity enables a rapid response to infection but may carry a cost for uninfected individuals, leading to the prediction that it will be favoured when infection rates are high. When we exposed populations of *Drosophila melanogaster* to intense parasitism by the parasitoid wasp *Leptopilina boulardi*, they evolved resistance by developing a more reactive cellular immune response. Using single-cell RNA sequencing, we found that immune-inducible genes had become constitutively upregulated. This was the result of resistant larvae differentiating precursors of specialized immune cells called lamellocytes that were previously only produced after infection. Therefore, populations evolved resistance by genetically hard-wiring an induced immune response to become constitutive.

## Main

A fundamental division within immune systems is between induced and constitutive immune defences. Constitutive immunity is always active, even in the absence of infection. For example, uninfected individuals produce natural antibodies^1^, immune cells and antimicrobial peptides (AMPs) ^2^. In contrast, induced immune responses are triggered after infection, which frequently involve amplifying existing constitutive factors. This can include immune cells proliferating and differentiating, and the massive upregulation of AMPs and cyotoxic molecules ^2^. Constitutive immunity is fast, allowing hosts to avoid becoming infected altogether or to rapidly clear infections. However, immune defences can be costly, diverting limited resources from other fitness-related traits ^3^ or damaging the host through immunopathology ^4^. While induced immune responses only incur these costs after infection, they are always present for constitutive immunity, even in the absence of infection. This trade-off between speed and cost is predicted to govern the allocation of host investment into constitutive and induced immunity ^5,6^. Theoretical models predict that the key parameter determining which type of defence should be favoured is the frequency that hosts are exposed to infection—more frequent exposure favours constitutive defences ^5^. In support of this, bacteria tend to evolve phage resistance by altering surface receptors when exposure is high and rely on induced CRISPR-Cas immunity when exposure is low ^7^. However, to our knowledge the key prediction that high rates of parasitism lead to induced immune responses becoming constitutively expressed has not been demonstrated.

To investigate these questions, we examined how the constitutive and induced cellular immune defences of *Drosophila melanogaster* evolve under high parasite pressure. Cellular immunity in *D. melanogaster* involves blood cells called hemocytes, which play the equivalent role to leukocytes in vertebrates. In uninfected larvae there are two morphological classes of hemocytes. Plasmatocytes constitute the majority of cells and have diverse functions including phagocytosis and AMP production ^8^. Crystal cells are a less abundant specialized cell type that produces the prophenoloxidase molecules PPO1 and PPO2, which are processed into enzymes required for the melanization of parasites ^9^. Infection can trigger an induced response, where these cell types proliferate, and cells called lamellocytes differentiate from plasmatocytes and prohemocytes in the lymph gland ^8^. Lamellocytes are large flat cells with a specialised role in killing large parasites like parasitic wasps (parasitoids). When a parasitoid injects its egg into *D. melanogaster* larvae, plasmatocytes adhere to the egg. Then, as lamellocytes differentiate they create additional cellular layers known as a ‘capsule’. Finally, the capsule is melanised by the activity of phenoloxidases produced in crystal cells and lamellocytes ^9^. The melanin physically encases the parasitoid and toxic by-products of the melanisation reaction likely contribute to parasite killing ^10^.

There is substantial genetic variation in susceptibility to parasitoid infection within *D. melanogaster* populations ^11^, and this is associated with differences in hemocyte composition ^12,13^. Here, we used a combination of experimental evolution with physiological assays and single-cell RNA sequencing to examine how a high rate of parasitism by the parasitoid wasp *L. boulardi* affects constitutive and induced immune defences.

### The evolution of resistance is associated with a more reactive immune response

To investigate the evolution of the cellular immune response, we allowed genetically diverse populations of *D. melanogaster* to evolve under intense levels of parasitism. We established an outbred population from 377 wild-caught *D. melanogaster* females, and used this to found six experimental populations that were maintained at population sizes of 200 flies. In three of these populations, larvae were infected every generation, and flies that survived by encapsulating and melanising the wasp were used to establish the next generation (High Parasitism populations). The other three populations were maintained in the same conditions but without infection (No Parasitism populations). In line with previous studies ^12,14^, the High Parasitism populations increased in resistance over the generations (Fig. 1A, Selection regime x Generation: χ^2^= 91.41, d.f.= 1, *p*<10^−15^). After 33 generations, a mean of 76% of flies survived infection in the High Parasitism populations, compared to 10% in the No Parasitism controls.

**Figure 1.**
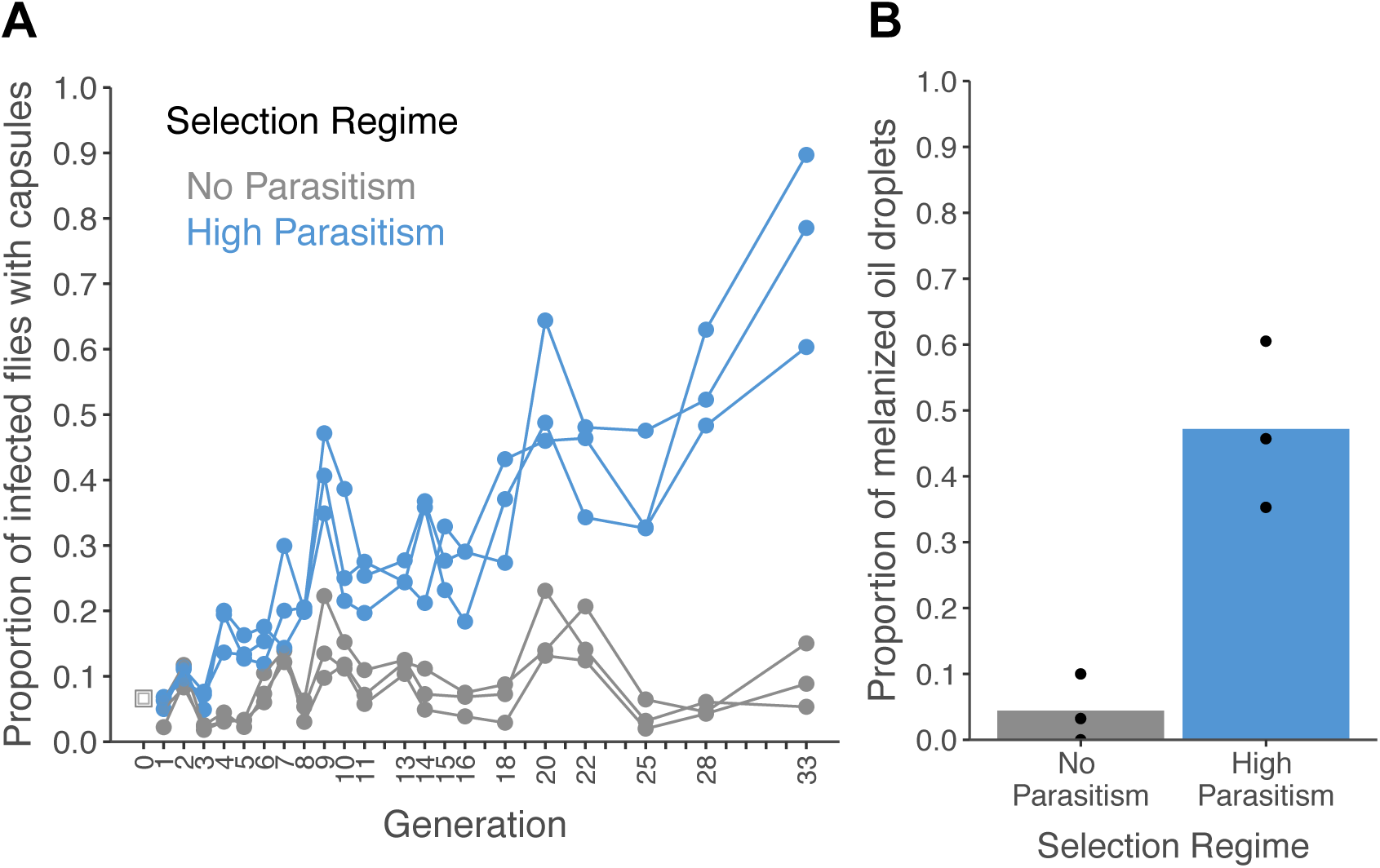
Encapsulation rate during selection for parasitoid resistance. (A) The proportion of infected larvae that gave rise to adult flies with visible capsules. An outbred population of *D. melanogaster* (source, grey square) was used to found six populations that were parasitized every generation with *L. boulardi* (blue) or maintained without infection (grey). (B) The proportion of larvae encapsulating oil droplets at generation 33. Dots represent the proportion of larvae in replicate populations and bars represent the mean per selection regime.

As populations evolved resistance, they melanised parasitoids at a higher rate (Fig. 1A). This could result from increased basal reactivity of the encapsulation immune response, or because the flies are evolving to escape the effects of immunosuppressive molecules produced by the parasite. To distinguish these possibilities, we measured the ability of each population to react to an inert object. When a small volume of paraffin oil is injected into the larval haemocoel, it remains as a sphere and can be encapsulated ^15^. Populations that evolved without parasitism show a very limited reaction to the oil droplets (Fig. 1B). In contrast, approximately half of larvae from selected populations encapsulate the oil droplet (Fig. 1B; Selection Regime: *F*_1,4_=24.53, *p*=0.008). Therefore, high levels of parasitism led to the cellular immune system evolving a more reactive encapsulation response.

### Constitutive upregulation of immune-induced genes

Underlying induced immune responses is a transcriptional response to infection, so we examined how the hemocyte transcriptome altered in populations that evolved under high levels of parasitism. We harvested hemocytes from *Drosophila* larvae and sequenced the transcriptomes of individual cells using the 10X Genomics platform. Initially, we pooled the data from scRNA-seq to investigate global changes in hemocyte gene expression. As expected, 48 hours post infection there was a strong transcriptional response to infection (Fig. 2A, X axis; Data S1). Strikingly similar transcriptional changes occurred in uninfected flies that had evolved under high rates of parasitism for 26 generations (Fig. 2A, Y axis; Data S1). Taking the 173 genes that had at least a 50% change in transcript levels after infection or selection, in every case the direction of change was the same after infection and adaptation to high parasitism (Fig. 2A; Data S1). We verified that the evolved response mimicked the induced response by using quantitative PCR (qPCR) to measure the expression of the lamellocyte marker *atilla* and the anti-parasitoid effector gene *PPO3* (Fig. S1). Again, we found that high rates of parasitism led to these immune-induced genes becoming constitutively upregulated. Therefore, as populations evolved resistance, the induced transcriptional response to infection has become genetically hard-wired and constitutive.

**Figure 2.**
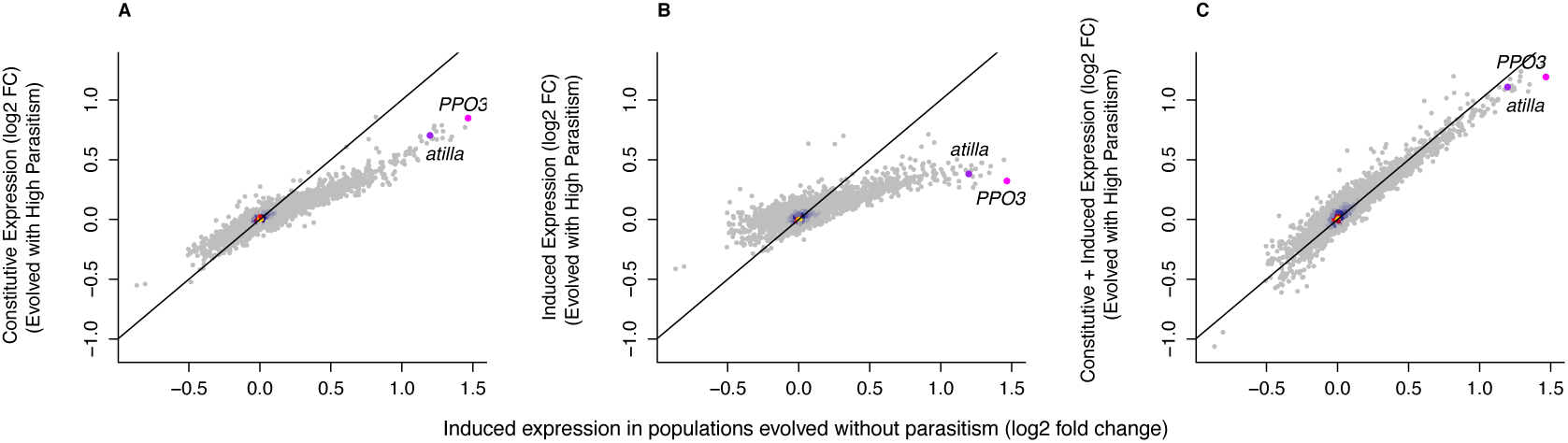
Changes in gene expression following selection for resistance and parasitoid infection. (A) Constitutive gene expression following 26 generations of selection for resistance to the parasitoid wasp *L. boulardi* (y-axis). (B) The induced change in gene expression following infection with *L. boulardi* in populations selected for resistance (y-axis). (C) The combined induced and constitutive change in gene expression following selection for resistance and infection (y-axis). All are compared to the change in gene expression following infection in populations maintained without parasitoid infection (x-axis). The black diagonal indicates the 1:1 line. The genes *atilla* and *PPO3* are highlighted. Colour represents the density of overlain points. Relative expression is expressed as log_2_(fold change).

We took the same approach to examine how the induced transcriptional response to infection changed as populations evolved resistance. We found that the transcriptional response to infection involved the same genes in the different populations (Fig. 2B). However, the magnitude of the changes in gene expression was substantially less in populations that had evolved under high parasite pressure (Fig. 2B). Therefore, the increase in constitutive gene expression has been matched by a decrease in induced expression. Again, this was verified by qPCR (Fig. S1). Combining the constitutive and induced changes in gene expression, the hemocyte transcriptome after infection is similar in the populations that evolved under high levels of parasitism and the controls (Fig. 2C). Therefore, an induced response has become genetically fixed, as opposed to selection leading to a greater total response.

### scRNA-seq reveals novel hemocyte types

The global changes in hemocyte transcriptomes could result from a change in the relative proportion of cell types in the hemocyte population, or from a shift in gene expression within existing cell types. The three classical *Drosophila* hemocyte types were originally defined morphologically, but emerging single cell transcriptomic data suggests that *Drosophila* hemocyte populations are more complex than this ^16–19^. Excluding cells that did not express the pan-hemocyte markers *Hemese* or *Serpent* ^20,21^, we grouped 19,344 larval hemocytes into nine clusters based on their transcriptomes (Fig. 3A). Neither infection nor selection leads to the appearance of new cell states (Fig. 3B).

**Figure 3.**
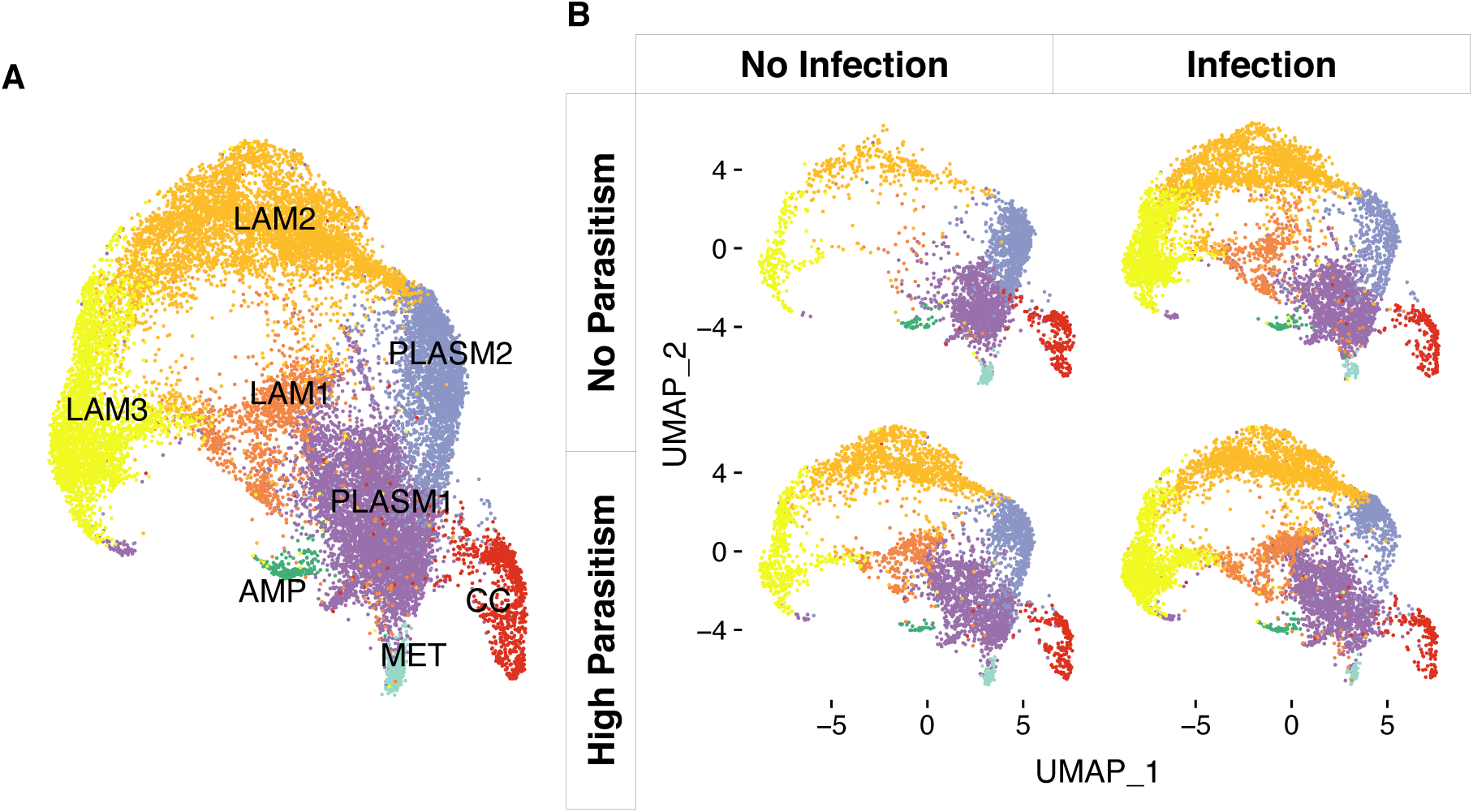
Cell states of *Drosophila melanogaster* larval hemocytes. Two-dimensional UMAP projections and cell state classification. (A) All 19,344 cells combined. PLASM: plasmatocytes; MET: metabolic plasmatocyte cluster; AMP: antimicrobial peptide plasmatocyte cluster; LAM: lamellocytes; CC: crystal cells. (B) Cells from populations that had undergone 26 generations of selection with no parasitism (top panels) or high parasitism (bottom panels), and were either uninfected (left panels) or 48 hours post infection (right panels). 64-333 million reads were sequenced in each of the eight 10X libraries that we generated (Data S2). The number of cells detected per library ranged from 787 to 3,463 (Fig. S6).

We identified two large clusters of plasmatocytes. The largest was PLASM1, which we classify as the cycling plasmatocyte state as it had the greatest number of replicating cells in the G2M or S phase (Fig. S2). The 5,712 cells in this cluster had high expression of genes associated with plasmatocytes (*Col4a1, Pxn, eater, Hml, NimC1*^21^) and actively replicating cells (*stg* ^22^, *polo*^23^and *scra*^24^) (Fig. S3). The next largest cluster, PLASM2, contained 2,874 plasmatocytes that were enriched for the genes encoding rRNA (Fig. S4) or involved in the ribosome biogenesis (Data S3). There were two smaller plasmatocyte clusters. The AMP cluster contained 196 cells with elevated expression of genes encoding antimicrobial and antifungal peptides, including *Drs, Mtk* and *CecA1* (Lemaitre and Hoffmann 2007, Fig. S4; Data S3). The MET cluster contained 283 cells with high expression of the heat shock proteins *Hsp24* and *Hsp27* and genes involved in organic acid metabolism (Fig. S4; Data S3). The crystal cells formed a single cluster of 906 cells (Fig. S6). Three clusters of cells, LAM 1, 2 and 3, had elevated expression of one or more lamellocyte marker genes ^21^ (Fig. S5) and contained 1,260, 5,104 and 3,000 cells respectively.

### Inducible cell states are present constitutively in resistant populations

Lamellocytes are key effector cells in the anti-parasitoid immune response. In populations that evolved without exposure to parasitoids, infection resulted in a large increase in the number of cells in all three lamellocyte clusters (LAM1-3, Fig. 3B and 4A). In populations that evolved under high levels of parasitism, this inducible response become constitutive, with uninfected larvae having 3-5 fold more cells in LAM1 and 2 (Fig. 4A). However, this was matched by a reduction in the induced response, so the final number of cells in these clusters after infection was similar regardless of whether the populations had adapted to high or low parasitism rates (Fig. 4A). These results held when replicate populations were analyzed separately (Fig. S7A, B) or when the number of cells in each library were down-sampled to match the library with the least number of cells (Fig. S7C). Therefore, high infection rates led to the inducible differentiation of specialised immune cells becoming genetically fixed.

**Figure 4.**
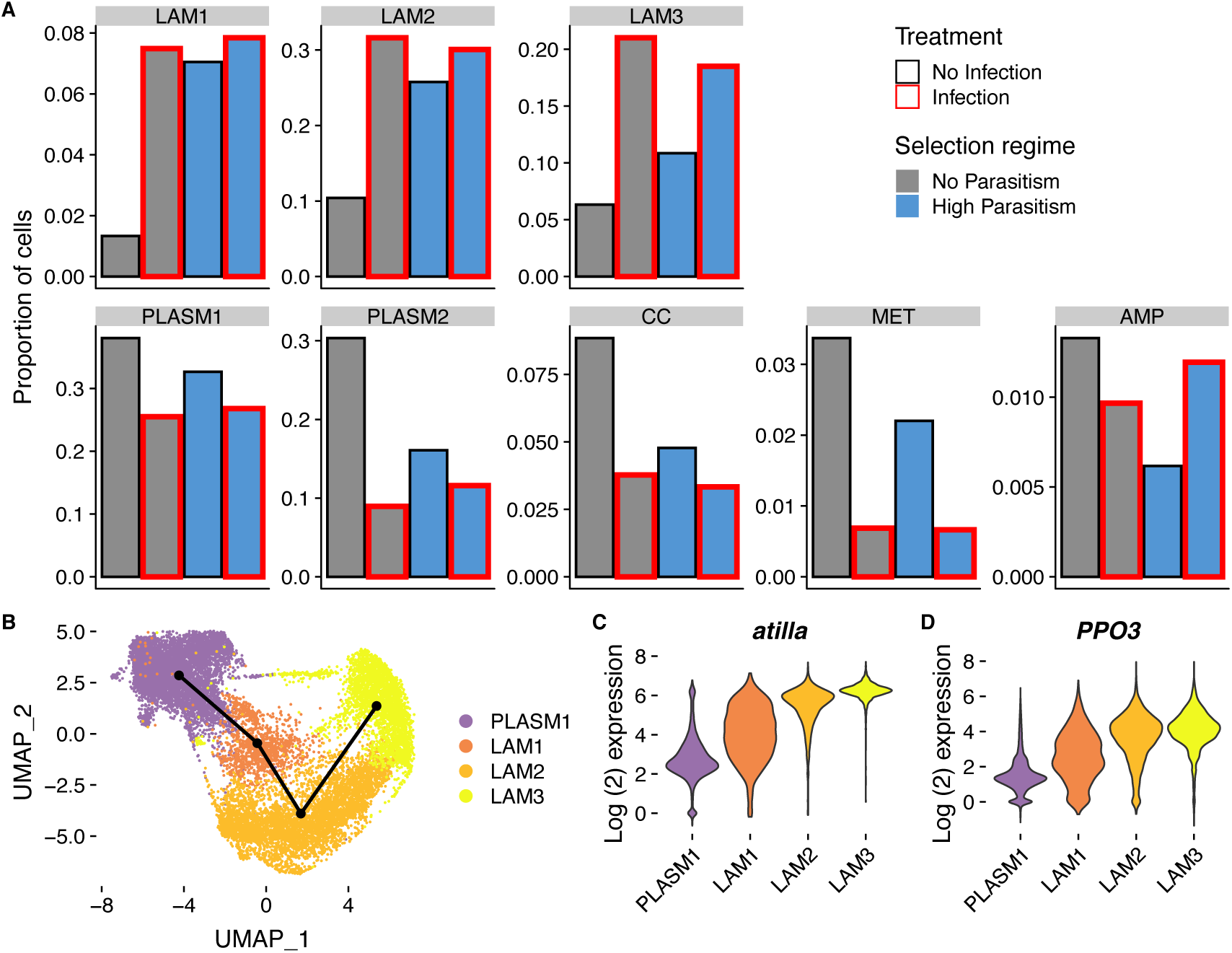
Changes in cell state following infection and adaptation to high rates of parasitism. (A) Proportion of different cell states after infection and selection. (B) Trajectory of lamellocyte differentiation. Plasmatocyte progenitors and lamellocytes were subclustered, the trajectories inferred from multi-dimensional principle components analysis, and the lineage projected onto the two-dimensional UMAP plot. (C-D) Log_2_ expression levels of *atilla* and *PPO3*. Relative expression levels were estimated from normalized and scaled unique molecular identifier counts.

To investigate the developmental origins of this change, we reconstructed the pathway by which plasmatocytes differentiate into mature lamellocytes. By subclustering the cells and ordering them along a pseudotemporal scale, we found a single lineage starting from the self-cycling plasmatocytes (PLASM1) and ending in LAM3 (Figs. 4B). Along this lineage there was a decrease in cell replication, with the largest fraction of cells in the G2M and S phase in PLASM1 and the largest fraction in G1 in LAM3 (Fig. S8). Importantly, controlling for the effects of cell cycle had a negligible impact on the clustering (Table S1) and the trajectory of lamellocyte differentiation (Fig. S9). The expression of the effector gene *PPO3*, the lamellocyte marker *atilla* and other marker genes all increased as cells progressed along this trajectory (Fig. 4C, 4D and S10, Data S4). On the other hand, genes involved in extracellular matrix organization—a house-keeping function of plasmatocytes ^25,26^—were down-regulated as cells differentiated (Data S5). Together, these results suggest that LAM1 cells are immature lamellocytes, while LAM3 cells are mature lamellocytes. Comparing our populations, the inducible production of LAM1 cells has become entirely constitutive in the populations evolved under high rates of parasitism, while much of the LAM3 response remains inducible. This suggests that high infection rates result in the cellular immune response becoming ‘primed’ by constitutively producing the precursors of the cells required for parasite killing.

### Morphological differentiation of immune cells remains an inducible response

Lamellocytes are classically defined morphologically as large flattened cells. As expected, the proportion of morphologically identifiable lamellocytes increased after infection (Figs 5A and Fig. S11A; Treatment: χ^2^= 48.14, d.f= 1, *p*=3.97×10^−12^). However, in contrast to our analysis of single cell transcriptomes, there was little difference between selection regimes in either the induced or constitutive proportion of these cells (Figs 5A and Fig. S11A; Selection regime x Treatment: χ^2^= 0.16, d.f= 1, *p*=0.69; Selection regime: χ^2^= 0.41, d.f= 1, *p*=0.52). This suggests that while the transcriptional state of cells has been constitutively activated, the change in cell morphology remains an inducible response. To test this hypothesis, we used microscopy to measure both cell morphology and, as a marker of the transcriptional state of the cells, *atilla* expression (Fig. S12). In line with our scRNAseq data, both infection and adaptation to high parasitism rates increases the proportion of *atilla* positive cells (Fig. 5C; Selection Regime x Treatment: *F*=16.3, d.f=1,8, *p*=0.004). However, among the *atilla* positive cells, infection had a large effect on cell size (Fig. 5C; Treatment: *F*=31.8, d.f.=1,7, *p*=0.0008), while selection had a small effect (Fig. 5C; Selection Regime: *F*=7.2, d.f.=1,7, *p*=0.03). Therefore, while selection has resulted in the constitutive transcriptional activation of cells, changes in cell morphology remain largely an inducible response.

**Figure 5.**
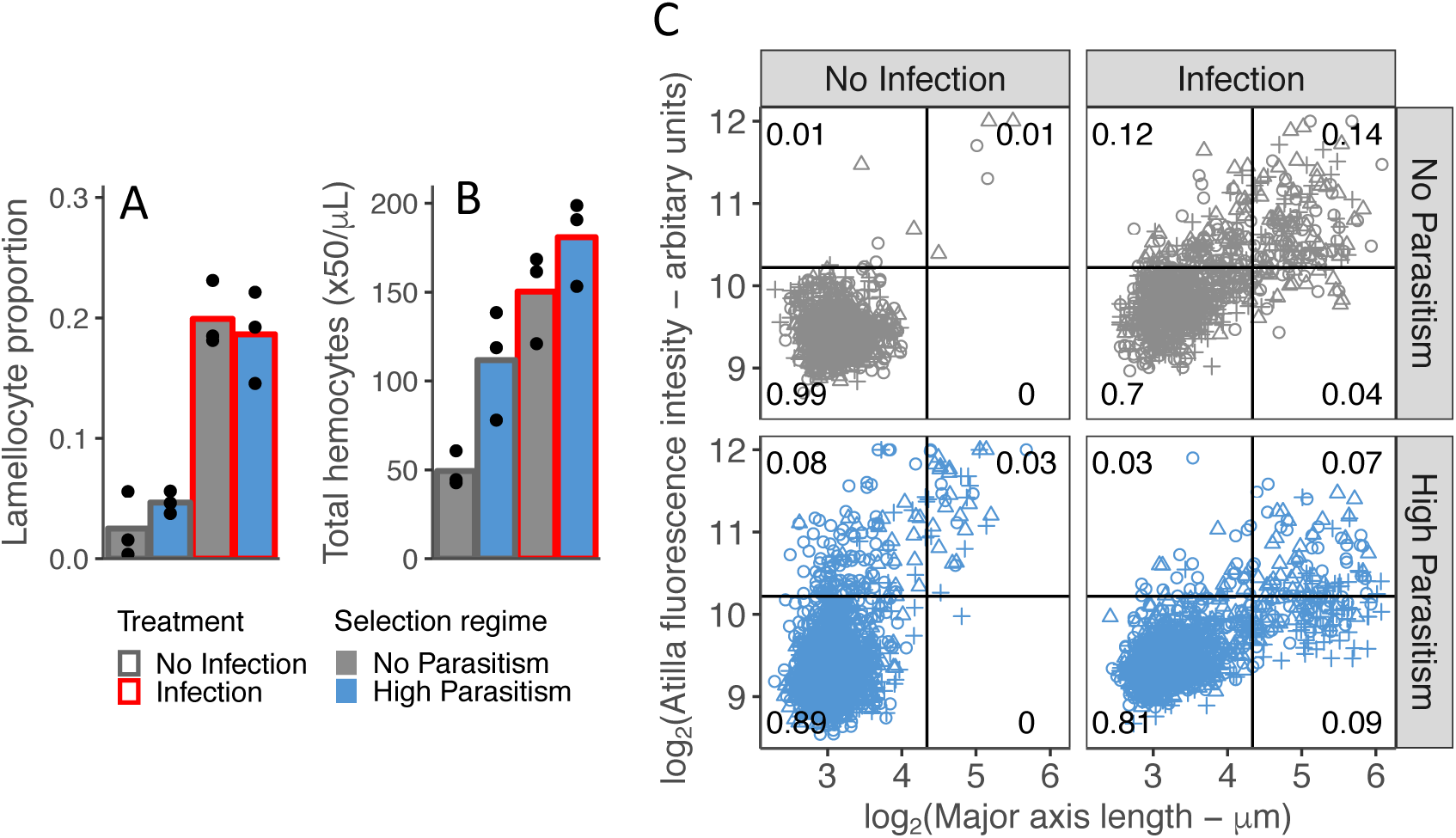
Changes in the morphology and number of cells following infection and adaptation to high parasitism rates. (A) The proportion hemocytes that were morphologically identified as lamellocytes and (B) the concentration of total circulating hemocytes in no infection conditions (grey outline) and 48 hours post infection (red outlines). Samples were collected from populations that evolved 34 generation with high parasitism (blue bars) or no parasitism (grey bars) (C) Cell size and *atilla* expression. The expression of *atilla* was measured as the fluorescence intensity resulting from immunohistochemistry staining. Cell size was measured along the major axis. Replicated populations from each selection regime are represented by different symbols and the proportion of cells in each quarter of the plot is shown.

In addition to the change in the transcriptional state of cells, we found that the populations maintained under strong parasite pressure evolved a constitutive increase in the absolute number of hemocytes in circulation (Fig. 5B and Fig. S11B, Selection regime: χ^2^= 22.37, d.f= 1, *p*=2.25×10^−6^). This supports results from previous selection experiments ^13,27^ and comparative analyses ^28^, and is another aspect of the induced response that has become constitutive.

## Conclusions

Our results provide support for the hypothesis that natural selection favours constitutive defences when infection is common ^5–7^ by demonstrating that an inducible immune response becomes constitutive when populations frequently encounter a parasite. The induced immune response to parasitoids relies on the differentiation of lamellocytes from embryonic-derived plasmatocytes, lymph gland prohemocytes ^8^ and lymph gland plasmatocytes ^18^. Evidence for immature lamellocytes has previously come from patterns of marker gene expression ^29,30^ and recent scRNAseq data ^17,19^. Our results show that *Drosophila* populations evolve to produce these cells constitutively if they are frequently exposed to infection. However, this change in the transcriptional state of cells was not accompanied by morphological differentiation. Genes involved in cytoskeleton dynamics such as *βTub60D* and *shot* are among the genes with the greatest increases in expression as lamellocytes differentiate (Fig. S10, Data S4) and these may be required for the final step of changing cell shape ^31^. Together, our results suggest populations that adapt to high parasitism rates have their cellular immune defences in a state of readiness, with the final step of cell differentiation remaining an inducible response.

Evolving resistance to parasitoid wasps is costly in *Drosophila* ^12,14,32^, so inducible immunity may be an adaptation that allows uninfected individuals to avoid this cost under low parasitism conditions. The physiological mechanism that causes the cost is unknown. It is possible that the differentiation of immature lamellocytes diverts resources from other fitness related traits. Lamellocyte differentiation is energy-demanding and results in the increased expression of glycolysis genes in hemocytes after infection ^3^. This cost may be borne by uninfected flies in our resistant populations, as we found that genes involved in trehalose uptake and catabolism ^33^, *Trehalase* and *Tret1-1*, are upregulated by both infection and selection (Fig. S13). Another possibility is that a more reactive cellular immune response may increase the effects of immunopathology.

After infection, whether the host or parasitoid survives is a race between the parasitoid development and the formation of a cellular capsule by the immune system ^34^. Constitutive differentiation of immature lamellocytes may allow lamellocytes to be produced more rapidly upon infection. Furthermore, cytotoxic molecules could be more readily available. This is supported by an increase in the constitutive expression of *PPO3* in populations adapted to high rates of parasitism. *PPO3* encodes a intracellular prophenoloxidase that, when active, is responsible for the melanization of lamellocytes ^9^. Along with the increase in the total number of hemocytes, the constitutive activation of the cellular immune response likely explains the greatly increased resistance of populations adapted to high rates of parasitism.

## Methods

### Artificial selection

An outcrossed *D. melanogaster* population was founded from the progeny of 377 females collected in July 2018 from banana and yeast traps set up in an allotment plot in Cambridge, UK (52°12’12.5”N 0°09’00.6”E). Single females were sorted into vials with cornmeal food (per 1200ml water: 13g agar, 105g dextrose, 105g maize, 23g yeast, 35ml Nipagin 10% w/v) to create isofemale lines. From the progeny of each isofemale line, 5 females and 5 males lines were sorted into a population cage to create the source population with 3770 flies. Flies from the source population were allowed to lay eggs overnight in 90mm agar plates (per 1500ml water: 45g agar, 50g dextrose, 500ml apple juice, 30ml Nipagin 10% w/v) spread with yeast paste (*Saccharomyces cerevisiae* – Sigma-Aldrich #YSC2). Eggs were removed from the agar plate with phosphate buffered saline (PBS) and a paintbrush, collected in 15ml centrifuge tubes and allowed to settle on the bottom of the tube. 500μl of egg solution were transferred into a 1.5ml microcentrifuge tube. 5μl of egg solution were transferred into plastic vials with cornmeal food and incubated at 25**°**C, in a 14 hours light/10 hours dark cycle and 70% humidity. 48 hours after egg transfer, a single female wasp was added into vials for infection treatments and allowed to infect larvae for 24 hours. Vials from infection and no infection treatments were incubated for 12 days in total. Flies from infection treatment with visible capsules were collected and randomly sorted into triplicate selection lines (NSRef1-3). Flies from no infection treatment were sorted into triplicate control populations (C1-3). Consecutive generations of selection were maintained with the same protocol described. Population sizes were maintained at 200 adult flies for all populations, at roughly 50% sex ratio.

### Estimation of encapsulation ratio

To determine encapsulation ability, 10 vials per line were prepared without infection and 20 vials with infection according to the selection protocol described above. Flies emerging from the no infection vials (control flies) were counted. Flies from infected vials were discriminated for the presence/absence of capsules by squishing anesthetized flies between two glass slides and observing under a dissecting microscope. Encapsulation ratio was calculated according to the formula:

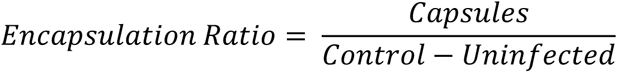

In the formula, Capsules represent the mean number of flies from infection vials with visible capsules and Uninfected represent the mean number of flies with no discernable capsule. Control represent the mean number of flies from vials with no infection. With this method we have an estimate of proportion of infected flies with successful encapsulation per line.

### Wasp maintenance

*Leptopilina boulardi* strain NSRef ^35^ was maintained in a very susceptible *D. melanogaster* outbreed population named CAMOP2. Cornmeal vials were prepared with 6μl of eggs as described above. Two female wasps and one male were added to each vial. Vials were incubated for 24 days at 25**°**C, in a 14 hours light/10 hours dark cycle and 70% humidity. Adult wasps were collected and maintained in cornmeal vials with a drop of honey.

### Oil injection

Egg solutions were prepared in 1.5ml microcentrifuge tubes as described above. 15μl of eggs were transferred into 50mm diameter plastic plates with cornmeal food. Plates were incubated for 72h at 25**°**C. Borosilicate glass 3.5” capillaries (Drummond Scientific Co. 3-000-203-G/X) were pulled to form thin needles in a needle puller (Narishige PC-10). The needle was backfilled with paraffin oil (Sigma-Aldrich #M5904) with a syringe and attached to a nanoinjector (Drummond Scientific Co. Nonoject II). Late 2^nd^ instar and early 3^rd^ instar larvae were carefully removed with forceps from cornmeal food plates and placed on filter paper, in groups of 20. Larvae were carefully injected with 4.6nl of oil. After injection, _dd_H_2_0 was added with a brush to remove the larvae and a total of 40 larvae were transferred into a cornmeal food vial and incubated at 25**°**C. After 48 hours larvae were removed with a 15% w/v sugar solution and transferred into _dd_H_2_O droplets on top of a plastic plate. Larvae were observed under a dissecting microscope and scored for the presence of melanization in the oil droplet.

### Hemocyte counts

To estimate hemocyte concentration, eggs were transferred into cornmeal vials following the artificial selection protocol described above and incubated at 25**°**C. In infection treatment vials, 3 female wasps were allowed to infect larvae for 3 hours, 48 hours post egg transfer. This protocol guarantees a high infection ratio ^36^ and reduces the time window for the treatment. Vials from all treatments were incubated for a total of 4 days. 3^rd^ instar larvae were collected from the food vials, washed in _dd_H_2_O, dried in filter paper and transferred to a multispot porcelain plate in groups of 10. Hemolymph was collected by tearing the larval cuticle from the ventral side. 2μl of hemolymph were rapidly collected and diluted in 8μl of Neutral Red (1.65g/L PBS – Sigma-Aldrich #N2889) to help visualize the cells. The diluted hemolymph was transferred into a Thoma chamber and the number of hemocytes in 0.1μl was counted. Plasmatocytes and lamellocytes were distinguished by morphology ^37^.

### Quantitative PCR

To analyze the expression of *PPO3* and *atilla*, RNA was extracted from hemocytes pooled from 50-70 larvae, 24h post infection. Larvae were cleaned in _dd_H_2_O, dried in filter paper and transferred into a well of a multispot porcelain plate with 100μl of PBS. Hemolymph was collected by tearing the ventral side of the larval cuticle. Samples were homogenized in 400μl TRIzol [Ambion #15596018] and kept at -80°C. For RNA extraction, samples were defrosted and centrifuged for 10min at 4°C at 12.000g. 160μl of supernatant was transferred into 1.5ml microcentrifuge tubes, 62.5μl of chloroform [Fisher Scientific #C/4920/08] was added, tubes were shaken for 15s and incubated for 3min. After a 10min centrifugation at 12,000g at 4°C, 66μl of the aqueous phase was transferred into a 1.5μl microcentrifuge tube, 156μl of isopropanol [Honeywell #33539] added and the solution thoroughly mixed. After 10min incubation samples were centrifuged for 10min at 12.000g at 4°C and the supernatant was removed. RNA was washed with 250μl 70% ethanol, centrifuged for 2min at 12.000g at 4°C. Ethanol was removed, samples dried, 20μl of nuclease free water [Ambion #AM9930] was added and samples incubated at 45°C for 10min. cDNA was prepared from RNA samples with GoScript reverse transcriptase (Promega) according to manufacture instructions. cDNA was diluted 1:10. Exonic primers for *D. melanogaster PPO3* and *atilla* were designed in NCBI Primer-BLAST online tool: (PPO3_qPCR_2_Fw: 5’-GATGTGGACCGGCCTAACAA-3’; PPO3_qPCR_2_Rev 5’-GATGCCCTTAGCGTCATCCA-3’; atilla_qPCR2_Fw: 5’-ACCCACCAAATATGCTGAAACA-3’; atilla_qPCR2_Rv: TTGATGGCCGATGCACTGT). The gene *RpL32* was used to normalize gene expression (RpL32_qPCR_F-d: 5’-TGCTAAGCTGTCGCACAAATGG-3’; RpL_qPCR_R-h 5’-TGCGCTTGTTCGATCCGTAAC-3’; ^38^. Sensifast Hi-Rox SyBr kit (Bioline) was used to perform the RT-qPCR on a StepOnePlus system (ThermoFisher Scientific). Each sample was duplicated (qPCR technical replica). The PCR cycle was 95°C for 2min followed by 40 cycles of 95°C for 5s, 60°C for 30s.

### Hemocyte immunohistochemistry

Larvae were prepared in the same conditions as for hemocyte counts, described above. Larvae were washed in _dd_H_2_O, dried on filter paper and transferred into a well of multispot porcelain plate with 25μl PBS. Hemolymph was collected by tearing the ventral cuticle with forceps and the hemocyte suspension was transferred into a reaction well of a microscope slide (Marienfeld, #1216330). Samples were incubated for 30min in a humid chamber, fixed for 20 min with 3.8% formaldehyde (Sigma-Aldrich #F8775) and washed 3 times with PBS. Cells were blocked and permeabilized with 0.01% Triton-X and 1% NGS diluted in PBS for 30 min and washed 3 times with PBS. Primary antibody against *atilla* (L1, 1:100, ^39^) was added with 1% NGS overnight at 4°C. Primary antibody was washed with PBS for 5 min 3 times and secondary antibody (Alexa Fluor 488, anti-mouse, 1:1000) was applied overnight at 4°C. Cells were washed 3 times with PBS and stained with Hoechst 33342 (Invitrogen #H3570, 1μg/ml) and phalloidin (Alexa Fluor 594, Invitrogen #A12381, dilution 1:2000) for 30min. After cells were washed 3 times with PBS, cover slides were mounted with Vectashield (Vector laboratories).

### Image acquisition and analysis

Images were acquired in a Leica DM6000 fluorescence microscope with a 40x objective. Images were analyzed with Fiji version 1.0 ^40^. Cells were identified based on actin and nuclear staining and manually outlined with the selection tool. Maximum intensity of *atilla* staining and length of the major axis was recorded for each cell. The threshold for staining intensity was obtained by measuring intensity of cells that were stained following the immunohistochemistry protocol (described above) but with no primary antibody. The threshold for cell size was obtained from a independent set of images where cells were measured and classified as lamellocytes.

### Statistical analysis

All statistical analyses were performed using R. Encapsulation proportions during artificial selection were arcsine transformed and analyzed with a general linear mixed effects model where selection treatment, generation and the interaction term were considered fixed factors and population a random variable. Oil melanization proportions were arcsine transformed and selection regime was used as an explanatory variable in a linear model. To analyze *atilla* and *PPO3* expression from qPCR results we used a mixed effects model with treatment, selection regime and the interaction term as fixed effects and population and target gene as random variables. Total hemocyte counts and the arcsine transformed lamellocyte proportions were analyzed with mixed effects models with treatment and selection regime as explanatory variables and population as a random variable. Proportions of cells considered atilla^+^ with immunohistochemistry staining were analyzed with a linear model with treatment, selection regime and the interaction term as explanatory variables. In all cases, significance was assessed with ANOVA.

### Hemocyte preparation for scRNAseq

Larvae from both selection regimes and infection treatments were obtained following the protocol for hemocyte counting, as described above. 80 to 140 larvae per biological replica were cleaned in PBS and dried on filter paper. Larvae were transferred into 200μl PBS and dissected by tearing the ventral cuticle. 200μl of hemocyte solution were collected and filtered (Flowmi cell strainer 70μm). The cell suspension was carefully transferred into a 15ml centrifuge tube with OptiPrep solution (Sigma-Aldrich D1556; 540μl OptiPrep and 1460μl PBS). Samples were centrifuged at 170g for 20 min in a swinging bucket centrifuge at 4°C with no brake. The first two layers of 100μl were then carefully removed and hemocytes were counted in a Thoma chamber. The sample fraction with more hemocytes was used for encapsulation in 10X single cell platform at Cancer Research UK Cambridge Institute. Library preparation was performed with standard protocols of single cell 3’ V2 chemistry 10X Genomics.

### Read alignments and cell detection

Demultiplexed FASTQ data files were aligned to a custom-built *D. melanogaster* reference obtained from FlyBase FB2018_02 Dmel Release 6.21 ^41^ and created using the Cell Ranger pipeline (https://support.10xgenomics.com/single-cell-gene-expression/software/overview/welcome, last accessed 23-03-2020). Less than 1,000 cells were detected in libraries generated from control and selected populations in one of the three replicates (replicate two). This replicate was dropped in further analysis. 25,128 cells were identified across the two remaining replicates. The entire sequence of data filtering and clustering is listed in Fig. S14. Most of the analyses were done in R versions 3.6 ^42^ and we used the Tidyverse, gridExtra and cowplot packages to manipulate and plot the data.

### Normalization, integration and dimensionality reduction

Seurat version 3.1.1 ^43^ was used to process and analyze the single cell count matrix. Unique molecular identifier (UMI) count matrices obtained from Cell Ranger were normalized using a scaling factor of 10,000 and log-normal transformation. 2,000 highly variable genes were discovered using the ‘vst’ method (Data S6). Count matrices from individual samples were integrated allowing for a maximum of 50 dimensions and 2,000 anchor features.

We identified 281 cells that expressed fewer than 250 features or more than 2,500 features and 5,387 cells that did not express either *He* or *Srp*, two pan-hemocyte markers ^21^. These cells were removed from the constituent samples. Following this, the count matrices were normalized, highly variable features were identified, and the samples were re-integrated.

### Mean fold change in gene expression

To combine data across cells to examine gene expression differences between treatments, we performed the data integration steps utilizing all possible anchors features in Seurat. In doing so, a total of 7,716 genes were used for integrating the sample count matrices (Data S6). Fold changes in expression between treatments were obtained from the output of *FindMarkers* with min.pct and logfc.threshold both set to 0. The fold change was converted to a log_2_ scale and plotted using the R package LSD.

### Clustering cells from scRNA-seq

Cells were clustered in Seurat. First, expression levels were scaled and percent mitochondria, number of features and UMI count per cell were regressed out. Then, we performed PCA and UMAP dimensionality reductions allowing for 50 dimensions. We did a first round of clustering using the ‘pca’ reduction and a clustering ‘resolution’ of 0.2. We identified markers upregulated in each cluster using the function *FindAllMarkers* with ‘min.pct’ and ‘logfc.threshold’ both set to 0.25 and tested for differential expression using MAST ^44^. We conduced enrichment analyses on upregulated cluster markers using the Flymine database and tool ^45^ with the 2,000 highly variable genes as the background list. We identified significantly enriched gene ontology categories ^46^ and pathways from the KEGG and Reactome ^47,48^ in each cluster.

Enrichment analyses from the first round of clustering indicated one cluster, comprising of 28 cells, was highly enriched for sperm-cell expressed genes (Data S7 - cluster 9). These cells were removed from the constituent sample count matrices and the data were re-integrated. In the second round of clustering, a ‘resolution’ of 0.3 was used and we discovered that cluster 9 (48 cells) and cluster 10 (41 cells) were enriched for genes associated with muscle and fat body tissues, respectively (Data S8). These cells were removed from the constituent sample count matrices and data were re-integrated. For round three of clustering, a ‘resolution’ of 0.3 was utilized. At this point, the 19,344 remaining cells were partitioned into nine clusters.

We identified the *D. melanogaster* orthologs of human cell cycle markers identified in Tirosh et al. ^49^ using the EBI database ^50^. The expression levels of these cell cycle genes were used to classify cells by phase using the function *CellCycleScoring* in Seurat. Regressing out the difference between the G2M and S phases had a minimal impact on the cell cluster classification for all but one cluster (Table S2). The greater discordance among the lamellocyte cluster CC-5 is due to the placement of cells differentiating along a single lineage into discrete clusters.

We tested if using all possible anchors rather than just the highly variable ones had an impact on data integration and clustering. Clustering the data set the 7,716 anchors using the same resolution (0.3) largely revealed the same cell clusters save for the appearance of two very small clusters comprising of 113 and 56 cells (cluster 9 and 10 - Table S3). Also, this data set showed that the proportion of lamellocytes (represented by clusters 1,2,4 and 5) increased following high parasitism and infection (Fig. S15) suggesting the choice of anchors did not result in erroneous patterns. When necessary, we converted the Flybase IDs to gene symbols using the org.Dm.eg.db database v.3.7 ^51^. The R package reshape2 ^52^ was used to estimate proportions of cells in each cluster.

### Subclustering lamellocytes

We performed subclustering on lamellocytes and their plasmatocyte progenitors to better resolve the trajectory of cell differentiation. Clusters 1, 2, 4 and 5 from round three clustering had enriched expression of described lamellocyte markers (*ItgaPS4, atilla, mys, PPO3* - ^21^) and, therefore, these cells were isolated for subclustering (Fig. S16). Cluster 0 from round three clustering was identified as the progenitor plasmatocyte population based on the expression of five plasmatocyte markers (*Col4a1, Pxn, eater, Hml, NimC1* - ^21^) and three cell cycle markers (*stg –* ^22^, *polo –* ^23^and *scra –* ^24^) (Fig. S17).

The presumed lamellocytes and plasmatocyte progenitor were subset from the constituent sample count matrices. The cell count matrices for each sample were normalized and 2,000 highly variable features were found. Following this, dimensionality reduction and round one of subclustering was performed using a ‘resolution’ parameter of 0.3. In round one of subclustering, we identified 79 cells that were significantly enriched for markers representing crystal cells (Fig. S18 & Data S9 - cluster 5). These cells were moved to the crystal cell cluster and we did a second round of subclustering on the remaining cells. We followed the same data integration and clustering steps but used a resolution of 0.2. The original cluster identities for lamellocytes and plasmatocyte progenitor cells were replaced with their new identities inferred from the subclustering.

### Trajectory inference

We used the software Slingshot ^53^ to conduct pseudotime analysis. We used all 50 PCA dimensions estimated from Seurat for trajectory inference and ensured lineages were not inferred past endpoints by setting stretch to 0. The cluster with the highest expression of the three cell cycle markers (*stg, polo* and *scra*) was set as starting cluster.

2000 highly variable genes were utilized for identifying lineage markers. A random forest model was fit to identify the genes best able to predict the pseudotime values using R tidymodels. The R package ‘Ranger’ was used for implementation of the random forest using mtry = 200, trees = 1400, and min_n = 15 and variable importance was ranked by Gini impurity.

### Cluster definitions

We performed pathway enrichment analyses on all non-lamellocyte clusters with the aim of defining them according to their inferred biological function. First, upregulated cluster markers were identified through pairwise differential expression tests. The cluster of interest was compared with PLASM1 using the function *FindMarkers* with ‘min.pct’ and ‘logfc.threshold’ both set to 0.25. Genes with significant upregulation in each cluster were identified using the MAST test statistic (Data S10) after performing Bonferroni correction on the obtained *P*-values. Markers upregulated in each cluster were used for pathway enrichment analysis. Markers for the plasmatocyte progenitor population were identified through comparisons to the cell types containing other plasmatocyte-type cells. These markers were used for pathway enrichment analyses using the approach described in previous sections.

## Acknowledgements

The Genomics Core Facility at Cancer Research UK Cambridge Institute, G. Raddi and S. Tattikota provided protocols for the scRNAseq.

## Author Contributions

AL, RA and FMJ designed the study and wrote the paper with input from the other authors. AL, JD and EG performed the *Drosophila* work. AL and JD generated the sequencing data. RA analysed the sequence data.

## Data and materials availability

Unprocessed single cell sequence reads were deposited in the Sequence Read Archive (accession: SRP256887, Bioproject: PRJNA625925). Cell count matrices for all detected genes, cluster identities and processed scRNA-seq results were deposited into Gene Expression Omnibus (accession: GSE148826). The R script used to analyze the scRNA-seq data is available on Github (Repository: dmel_scRNA_hemocyte).

