## supplementary material for "Constitutive activation of cellular immunity underlies the evolution of resistance to infection"

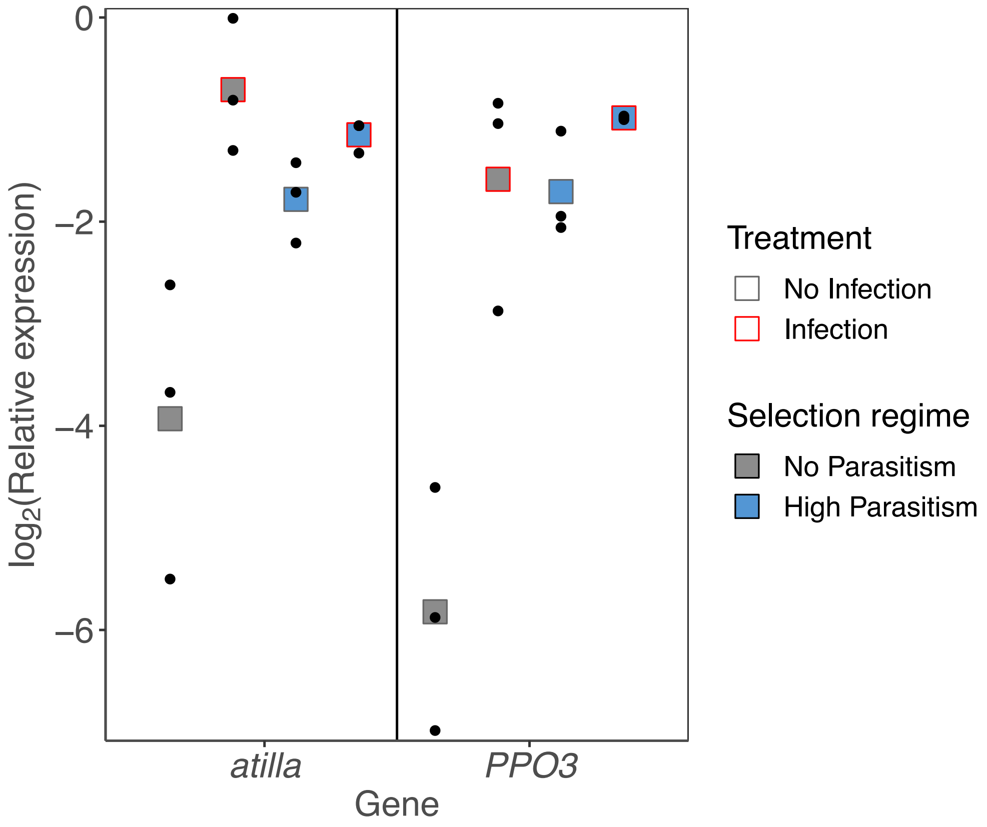


**Fig. S1 Gene expression induction in hemocytes after infection** *atilla* and *PPO3* expression normalized by the housekeeping gene *RpL32* in populations evolved under no parasitism (grey) or high parasitism (blue). Hemocyte samples for RNA extraction were collected from 50-70 larvae 24 hours post infection (red outlines) and from larvae with no infection (grey outline). Each dot represents the average ΔCt of two biological replicas per population and squares represents the average per selection regime. Selection regime x Treatment: $\chi 2$= 22.25, d.f= 1, *p*=2.39x10-6

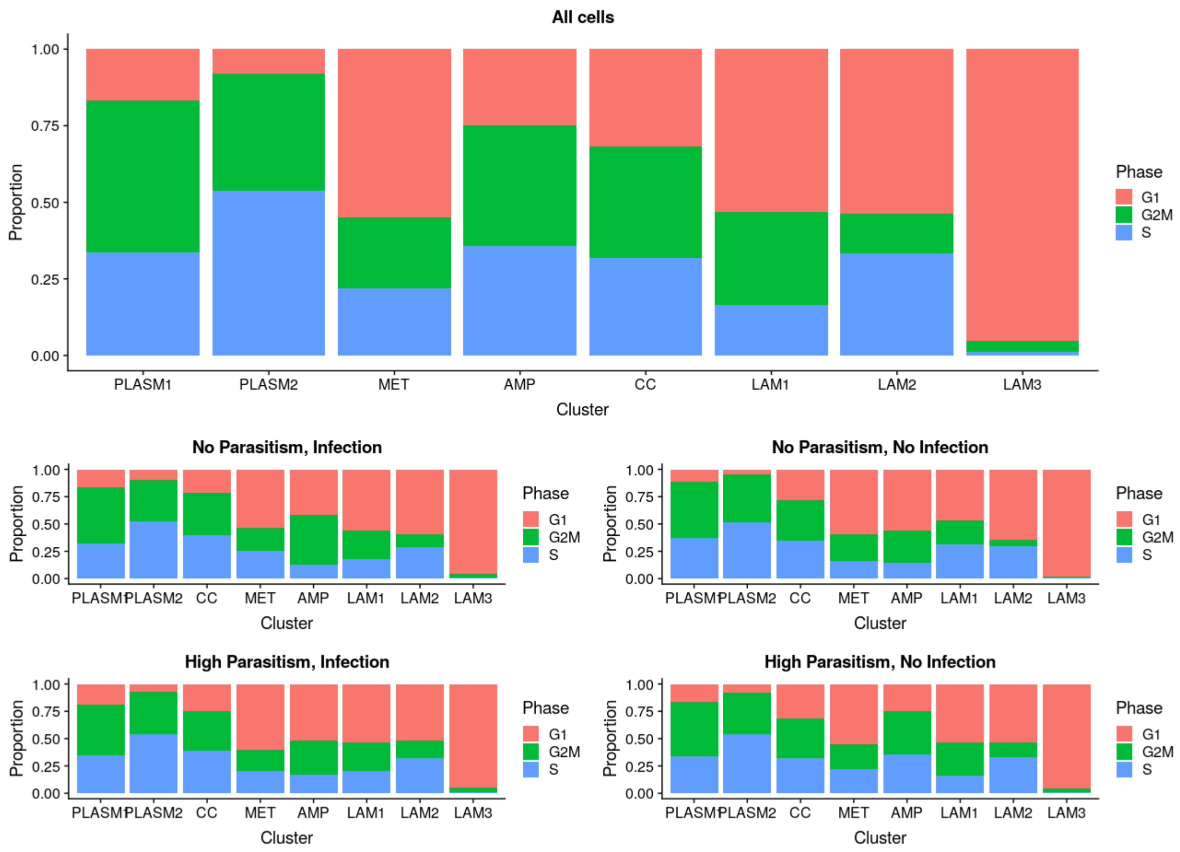


**Fig. S2.** Proportions of cells in G1, G2M and S phase in the plasmatocyte, lamellocyte and crystal cell clusters.

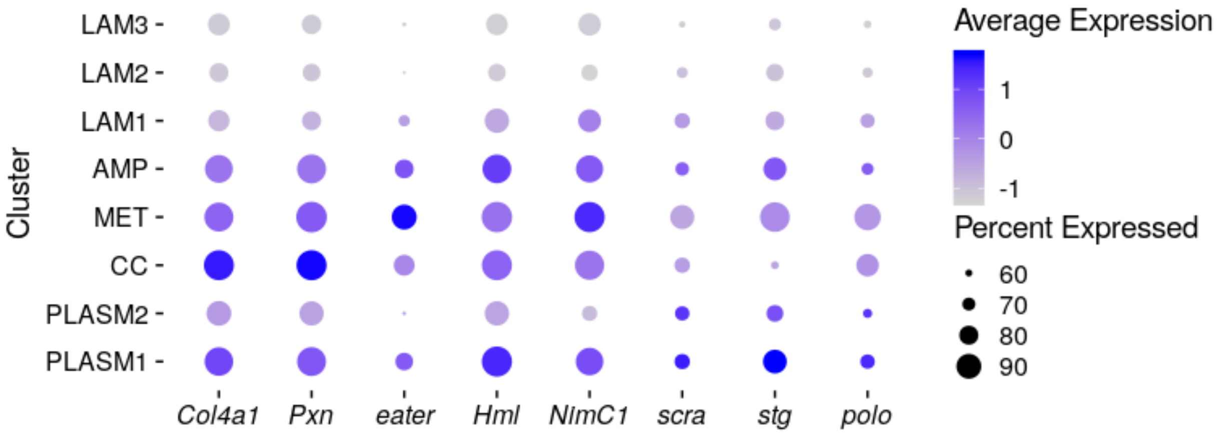


**Fig. S3.** Relative level of expression (log $_e$) of plasmatocyte and cell cycle marker genes. Percent of cells expressing genes are indicated by circle size.

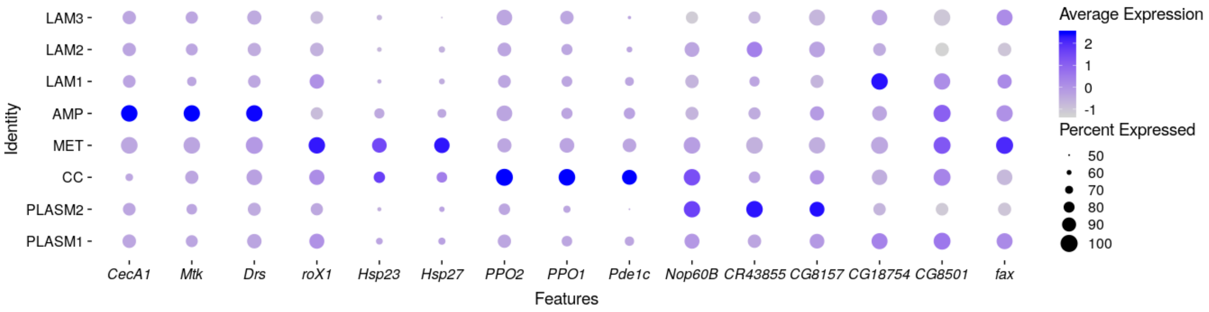


**Fig. S4.** Relative level of expression (log $_e$) of marker genes for plasmatocyte and crystal cell clusters. Percent of cells expressing genes are indicated by circle size.

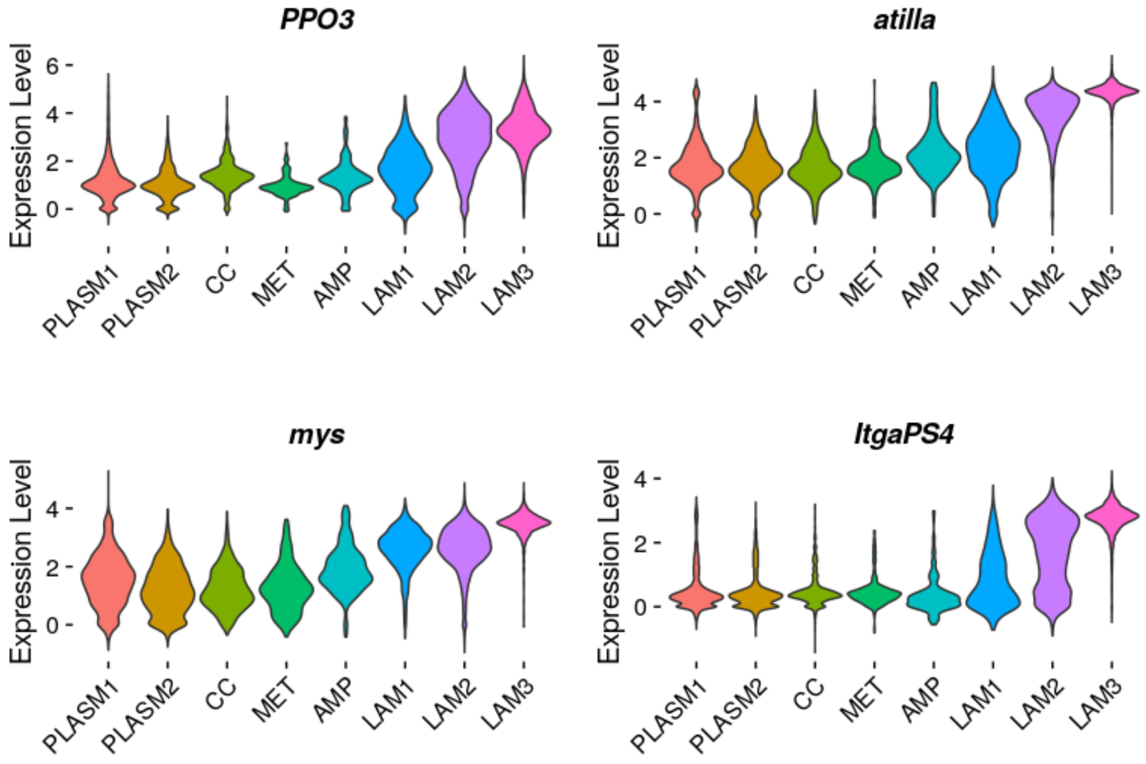


**Fig. S5.** Distribution of $\log_e$ gene expression levels for described lamellocyte marker genes grouped by cluster identities.

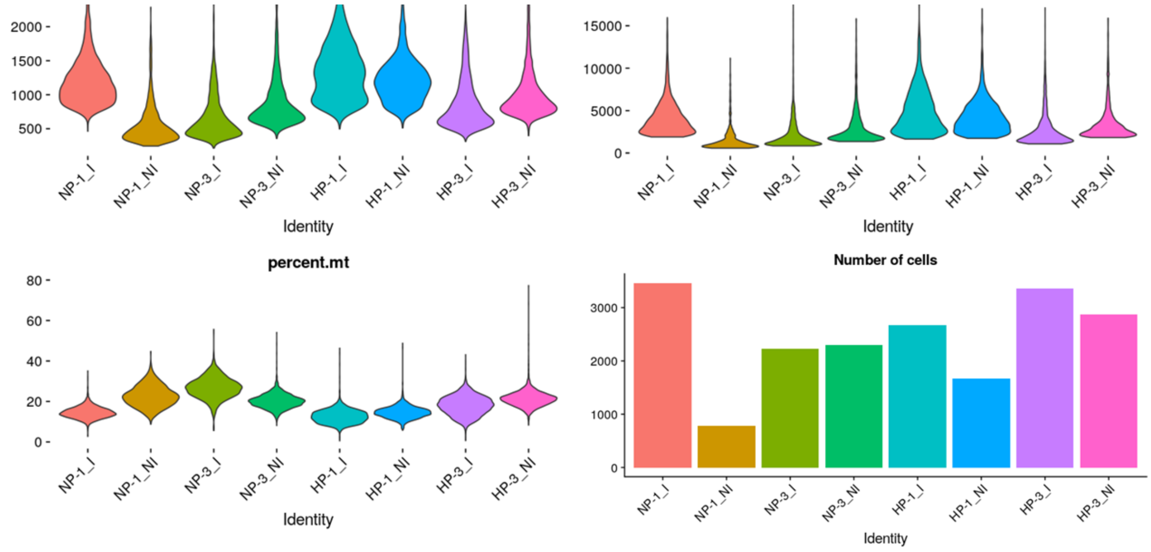


**Fig. S6.** Number of cells and genes detected, unique molecular identifier count and percent mitochondrial gene expression per cell grouped by sample. HP: high parasitism, NP: No parasitism, I: Infection and NI: no infection. Numbers following HP or NP correspond to replicate.

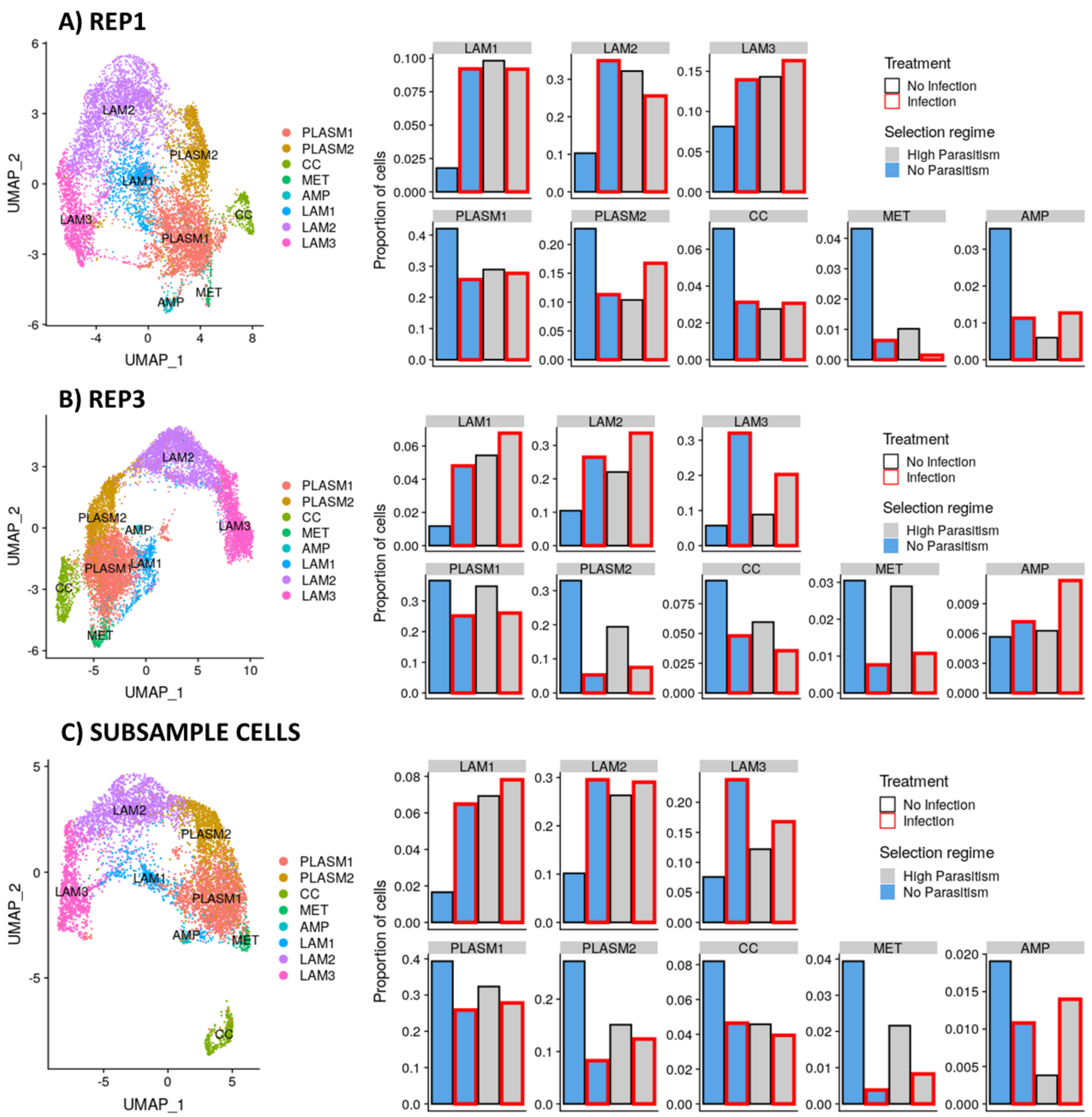


**Fig. S7.** Impact of cell number and library-specific differences on data integration, cell cluster classification and estimating cluster proportions. A) Clustering 8,596 cells from replicate one, B) clustering 10,784 cells from replicate three and C) clustering 6,296 cells where 787 cells were randomly subsampled without replacement in each of eight libraries used for data integration.

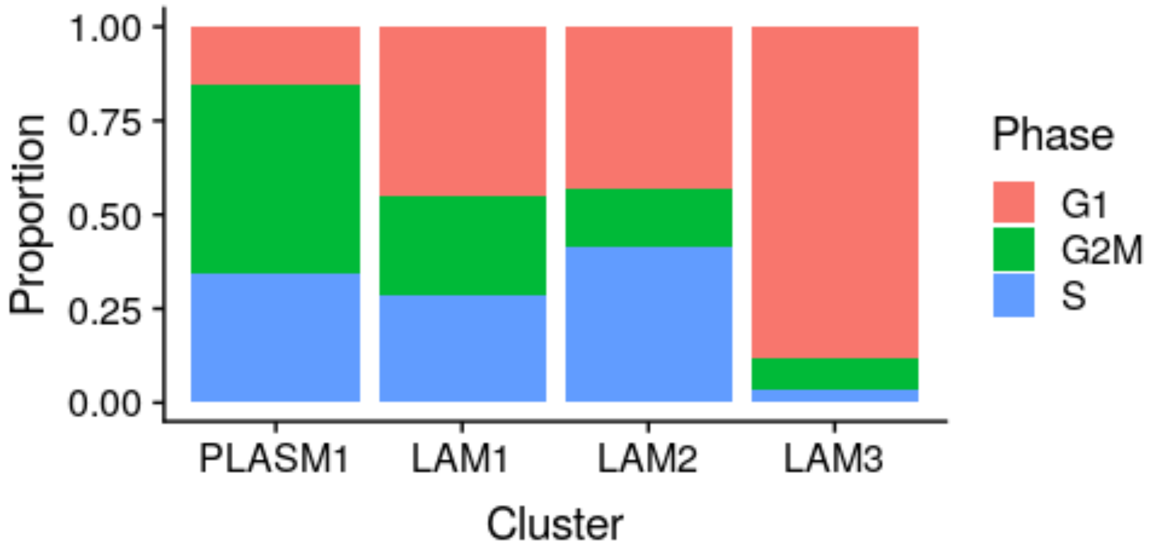


**Fig. S8.** Proportions of cells in G1, G2M and S phase in the plasmatocyte progenitor and lamellocyte clusters following subclustering.

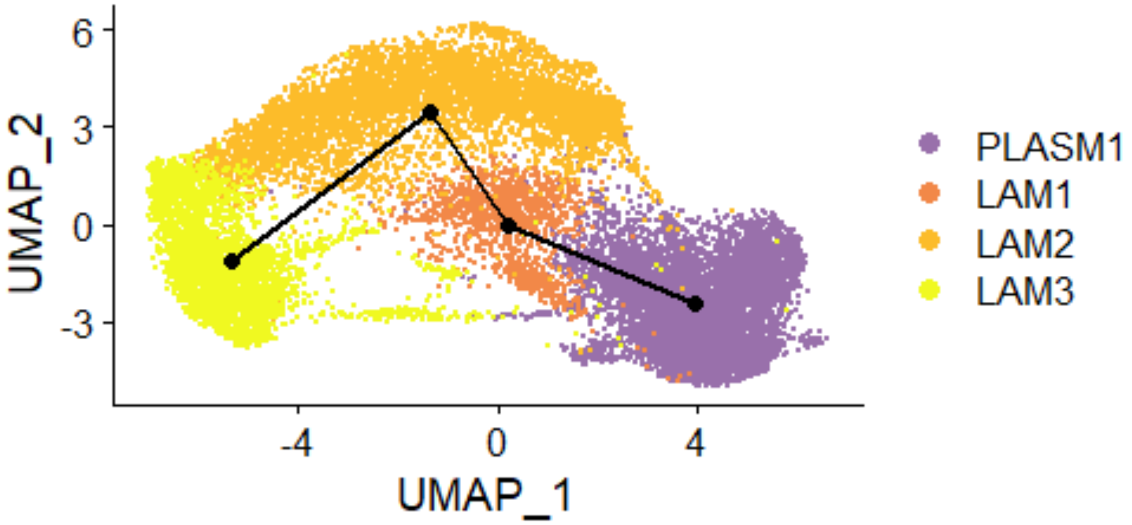


**Fig. S9.** Trajectory of lamellocyte differentiation inferred following cell cycle correction and regressing out the difference between the G2M and S phases. Trajectories were inferred from multi-dimensional principle components analysis, and the lineage projected onto the two-dimensional UMAP plot.

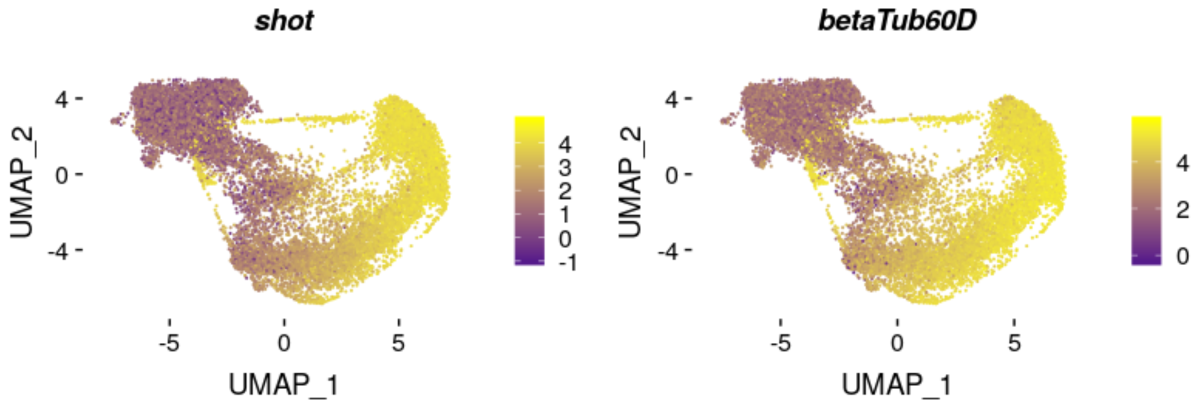


**Fig. S10.** $\log_e$ expression levels of *betaTub60D* and *shot*. Relative expression levels were estimated from normalized and scaled unique molecular identifier counts.

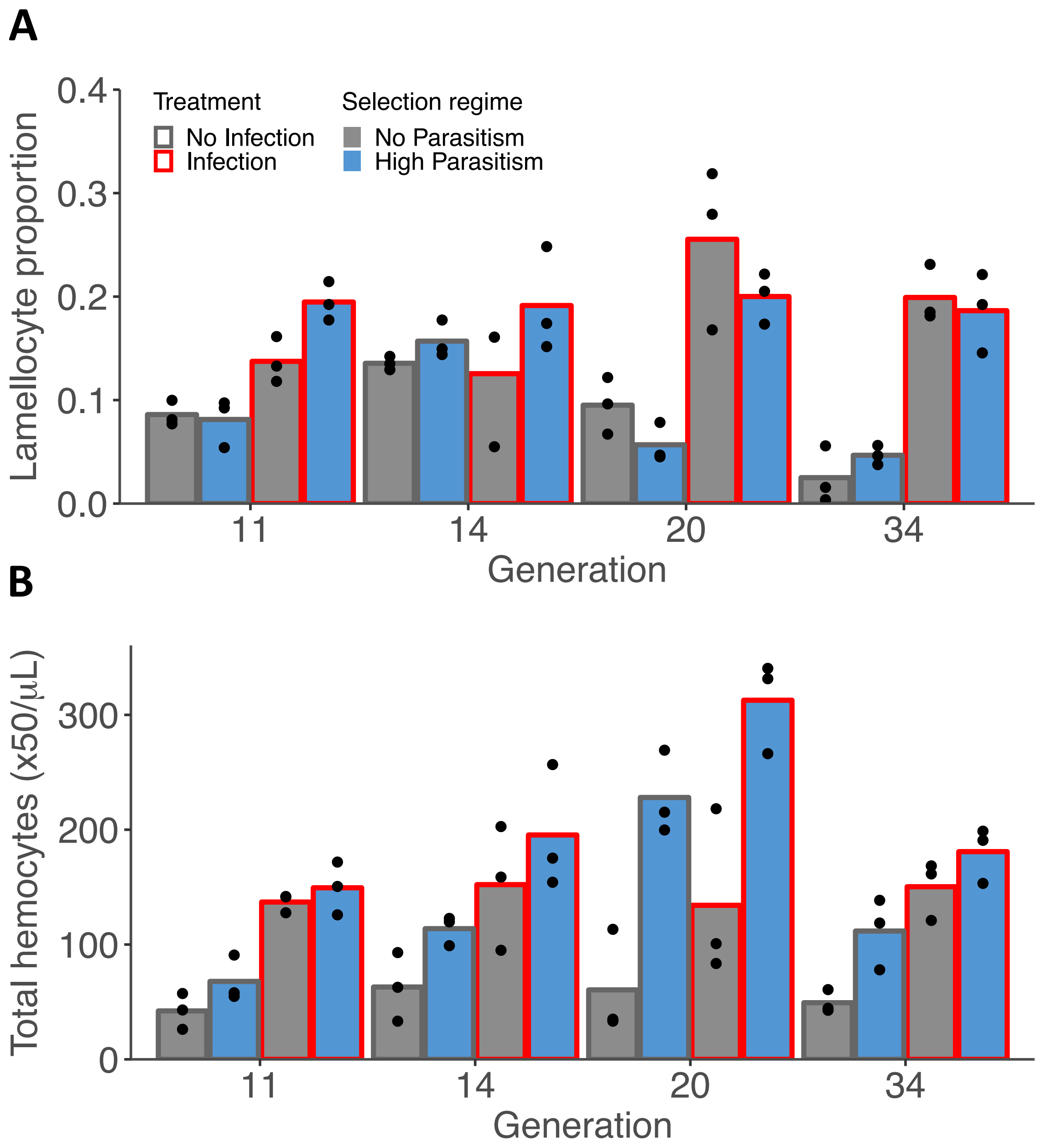


**Fig. S11. Changes in number of circulating hemocytes during artificial selection**
Concentration of total circulating hemocytes (A) and proportion of lamellocytes (B) in population evolved with no parasitism (grey bars) and with high parasitism (blue bars). Samples were collected as 3rd instar larvae in homeostasis (grey outline) or 48 hours post infection (red outline). Each dot represents the mean counts of each population calculated from 4-10 replicas and bar height represents the mean of the triplicate lines. Generation 34 is represented in Figure 5.

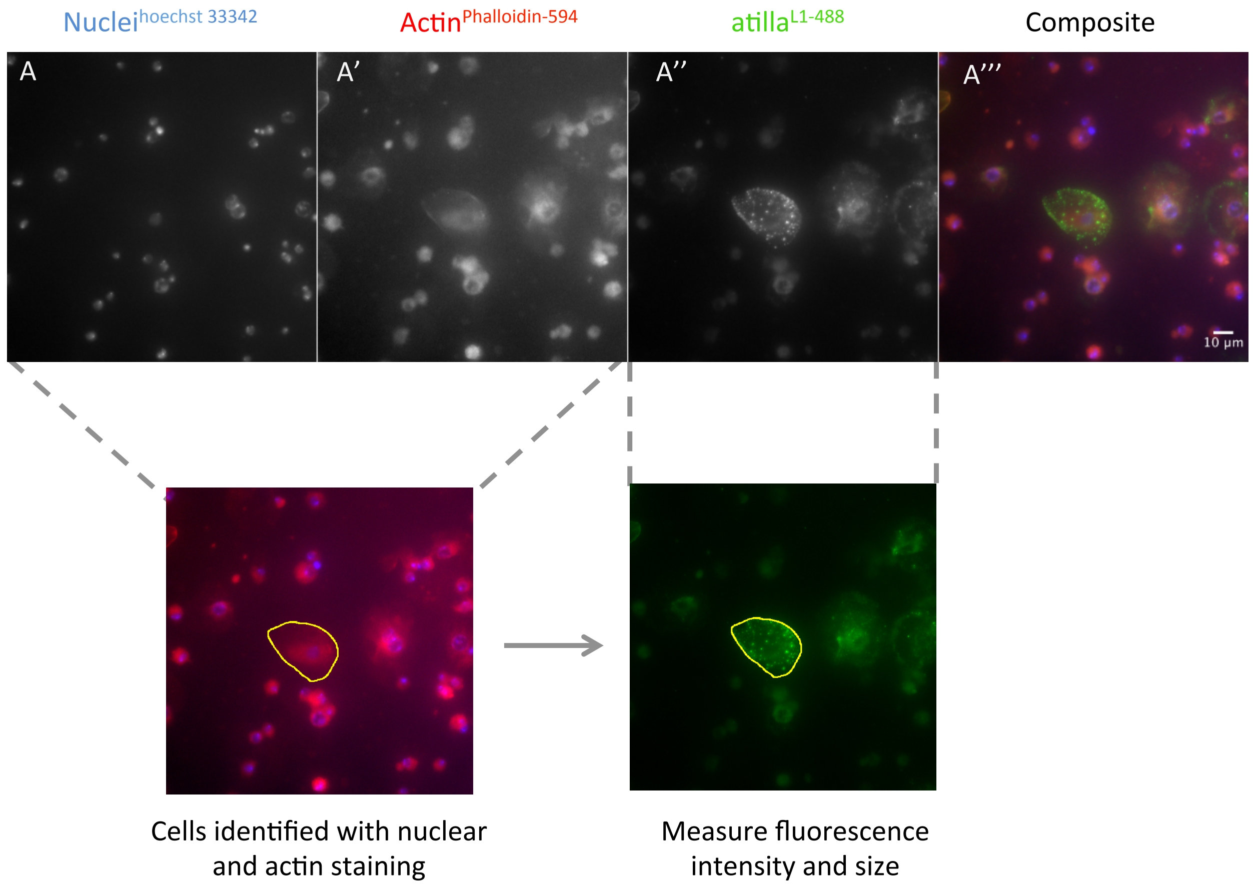


**Fig. S12. Hemocyte size and atilla staining intensity.** Hemocytes were stained for nuclei (A, hoechst 33342), Actin (A’, Phalloidin- Alexa594) and atilla (A’’, L1 antibody + Alexa488). Composite images were created for phallodin and hoechst staining to identify cells with selection tool (yellow line) and fluorescence intensity was measured in L1 staining.

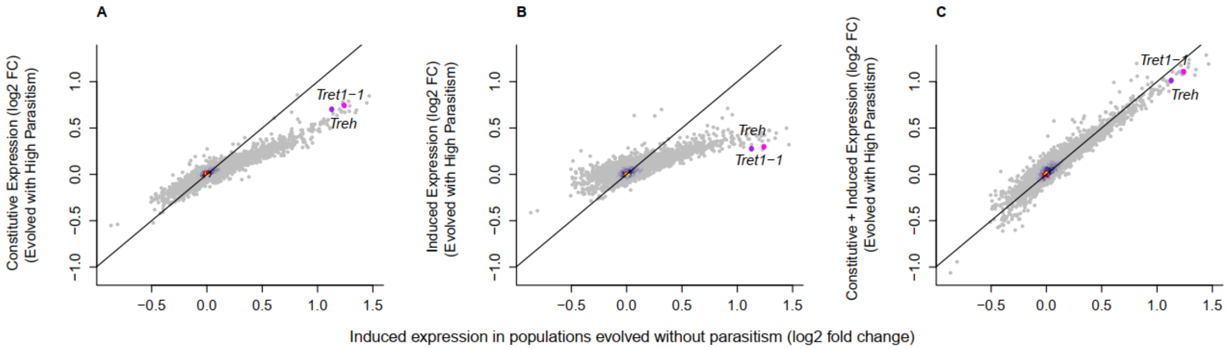


**Fig. S13. Changes in *Trehalase* and *Tret1-1* expression following selection for resistance and parasitoid infection.** (A) Constitutive gene expression following 26 generations of selection for resistance to the parasitoid wasp *L. boulardi* (y-axis). (B) The induced change in gene expression following infection with *L. boulardi* in populations selected for resistance (y-axis). (C) The combined induced and constitutive change in gene expression following selection for resistance and infection (y-axis). All are compared to the change in gene expression following infection in populations maintained without parasitoid infection (x-axis). The black diagonal indicates the 1:1 line. The genes *Trehalase* and *Tret1-1* are highlighted. Colour represents the density of overlain points.

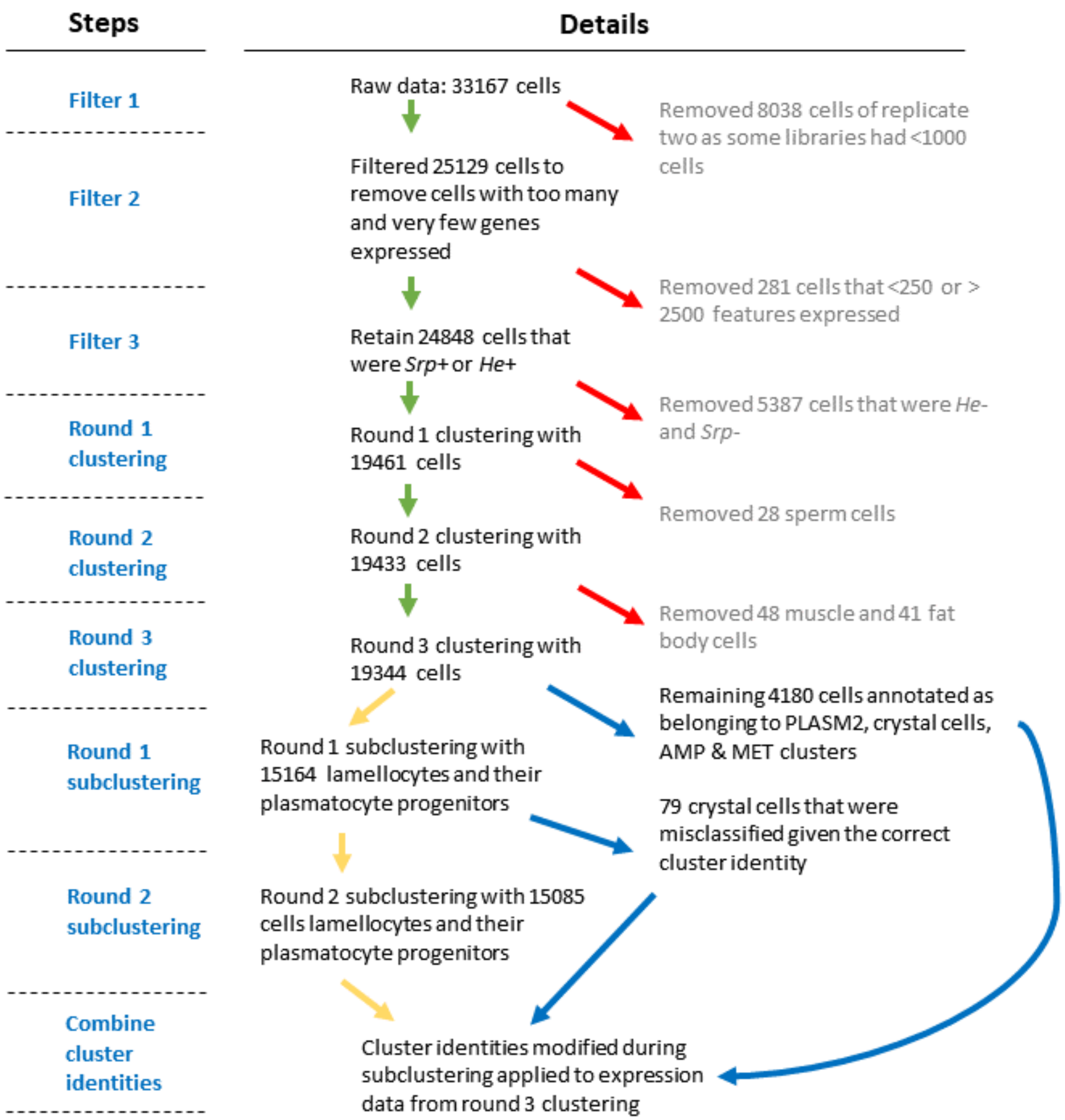


**Fig S14.** Flowchart of data filtering and clustering steps. Only cells expressing >250 features and < 2500 features and cells that were either *He*+ or *Srp*+ were kept. Cells expressing high levels of sperm-cell, muscle or fat body marker genes were identified during three rounds of clustering and removed. 19,344 cells remained after filtering. Two rounds of subclustering identified plasmatocyte progenitor and lamellocyte clusters and these identities replaced those attained from round three clustering for relevant cells.

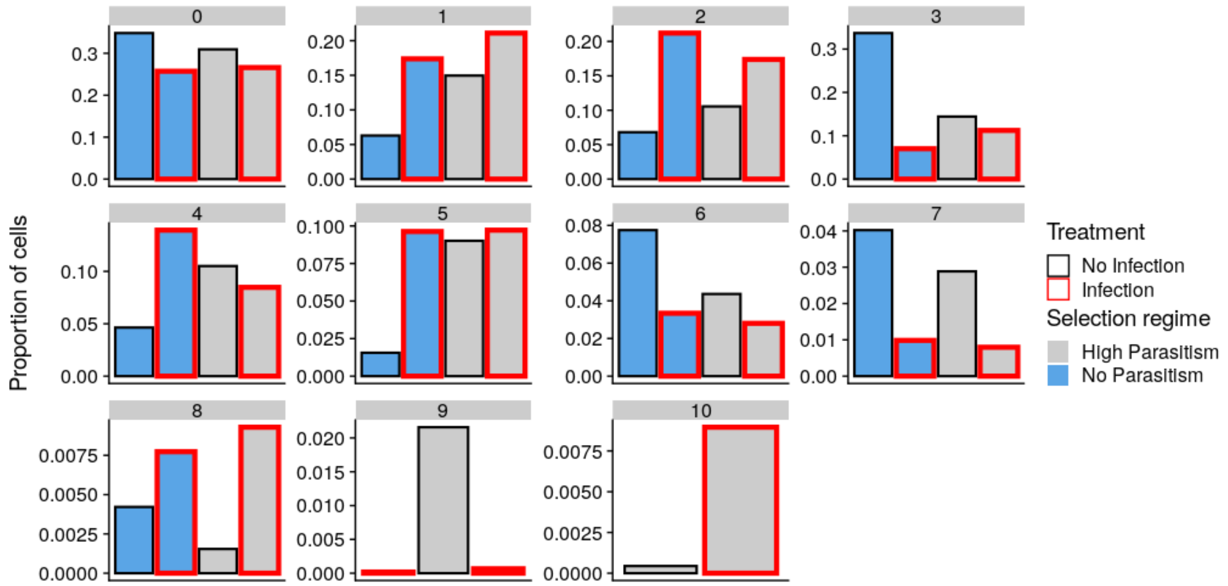


**Fig S15.** Proportion of different cell states after infection and selection following data integration with 7,716 anchors, the maximum number that can be detected.

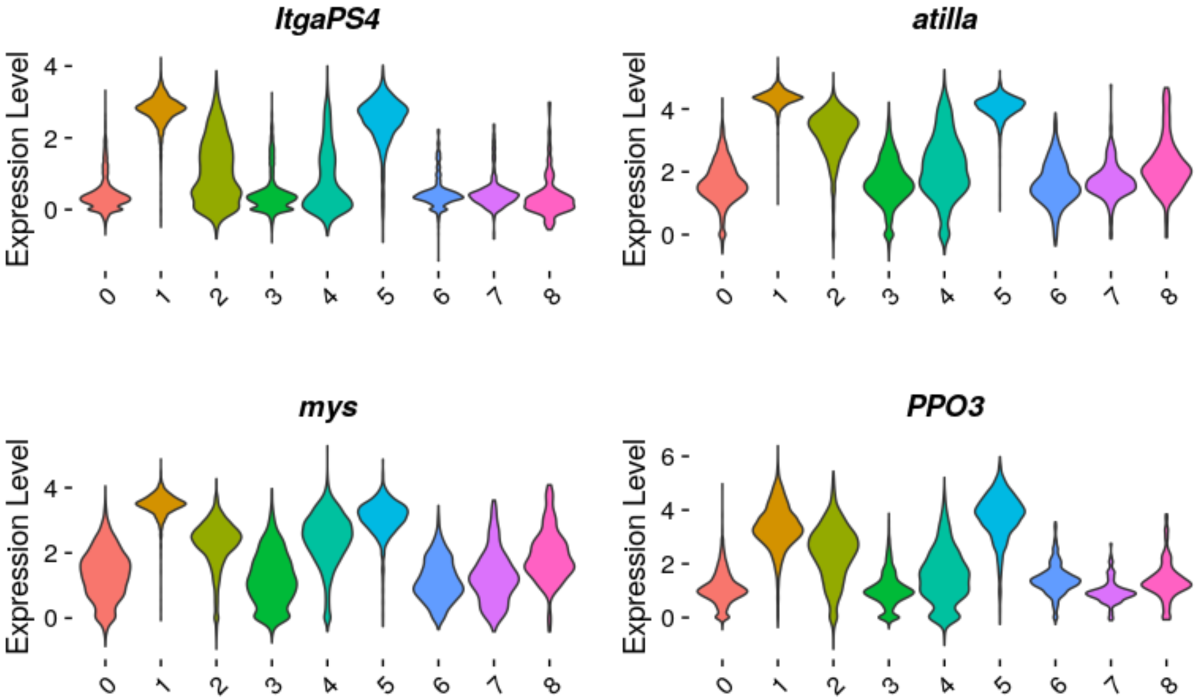


**Fig S16.** Distribution of gene expression levels ($\log_e$) of described lamellocyte marker genes, for clusters identified from round three clustering.

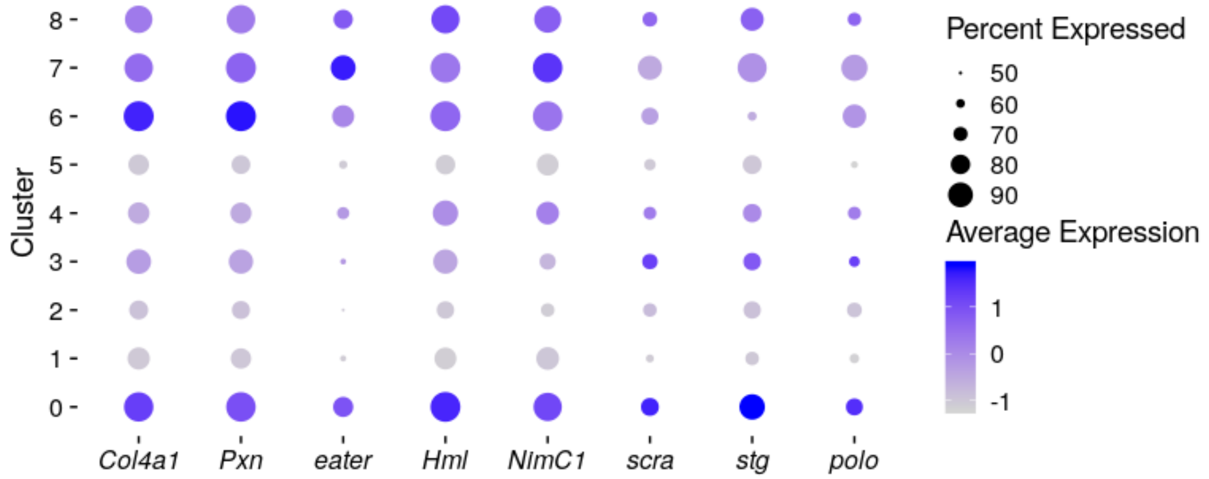


**Fig S17.** Relative level of expression (log $_e$) of plasmatocyte and cell cycle marker genes, for clusters identified from round three clustering. Percent of cells expressing genes are indicated by circle size.

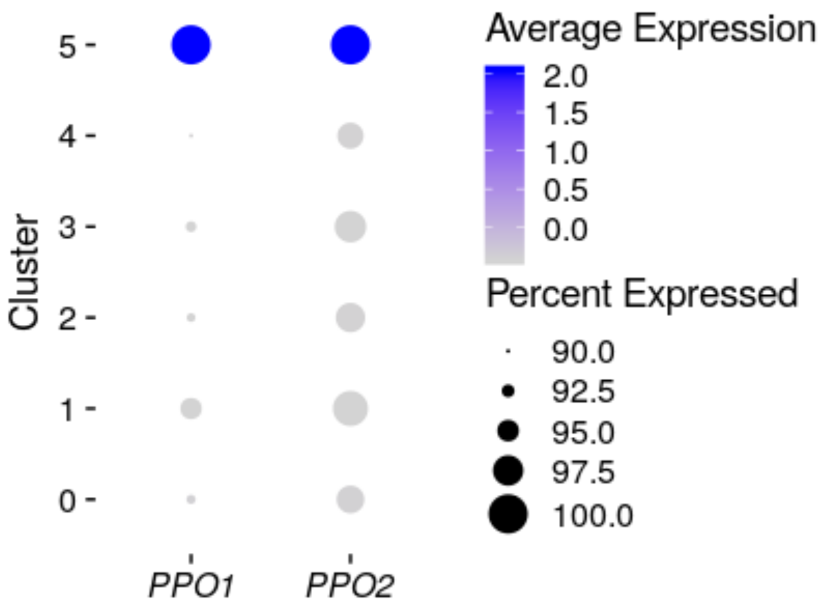


**Fig S18.** Relative level of expression (log $_e$) of two known crystal cell markers from round one of subclustering, using lamellocytes and their plasmatocyte progenitors. Percent of cells expressing genes are indicated by circle size.

| Cluster identity | NOCC-PLASM1 | NOCC-LAM1 | NOCC-LAM2 | NOCC-LAM3 |
|---|---|---|---|---|
| **CC-PLASM1** | 5662 | 20 | 33 | 6 |
| **CC-LAM1** | 88 | 1111 | 54 | 7 |
| **CC-LAM2** | 51 | 40 | 4954 | 59 |
| **CC-LAM3** | 2 | 0 | 25 | 2973 |

CC: Cell cycle effects corrected, NPCC: no cell cycle correction applied

**Table S1.** Comparison of subclustering classification of lamellocytes and their plasmatocyte progenitor cells with and without corrections for cell cycle effects.

| Cluster identity | CC-0 | CC-1 | CC-2 | CC-3 | CC-4 | CC-5 | CC-6 | CC-7 | CC-8 |
|---|---|---|---|---|---|---|---|---|---|
| **PLASM1** | 5340 | 50 | 83 | 143 | 67 | 36 | 0 | 1 | 1 |
| **PLASM2** | 255 | 12 | 2593 | 12 | 0 | 0 | 2 | 0 | 0 |
| **CC** | 80 | 12 | 1 | 1 | 5 | 2 | 805 | 0 | 0 |
| **MET** | 17 | 0 | 0 | 0 | 0 | 0 | 0 | 266 | 0 |
| **AMP** | 17 | 0 | 0 | 0 | 1 | 0 | 0 | 0 | 178 |
| **LAM1** | 270 | 47 | 0 | 1 | 942 | 0 | 0 | 0 | 0 |
| **LAM2** | 47 | 3894 | 267 | 10 | 187 | 698 | 0 | 0 | 1 |
| **LAM3** | 13 | 20 | 0 | 1976 | 267 | 723 | 0 | 0 | 1 |

CC: Cell cycle effects corrected

**Table S2.** Comparison of cell cluster classification of 19,344 hemocytes with and without corrections for cell cycle effects.

| Cluster identity | 0 | 1 | 2 | 3 | 4 | 5 | 6 | 7 | 8 | 9 | 10 |
|---|---|---|---|---|---|---|---|---|---|---|---|
| **PLASM1** | 4821 | 41 | 159 | 291 | 42 | 188 | 13 | 100 | 11 | 55 | 0 |
| **PLASM2** | 362 | 109 | 3 | 2338 | 0 | 7 | 7 | 2 | 1 | 45 | 0 |
| **CC** | 77 | 14 | 2 | 23 | 6 | 4 | 776 | 3 | 0 | 1 | 0 |
| **MET** | 32 | 0 | 0 | 5 | 0 | 1 | 0 | 245 | 0 | 0 | 0 |
| **AMP** | 61 | 6 | 1 | 5 | 1 | 15 | 0 | 1 | 105 | 1 | 0 |
| **LAM1** | 145 | 75 | 3 | 7 | 7 | 1015 | 0 | 6 | 2 | 0 | 0 |
| **LAM2** | 43 | 2886 | 127 | 100 | 1758 | 186 | 0 | 1 | 1 | 0 | 2 |
| **LAM3** | 6 | 3 | 2649 | 1 | 110 | 175 | 0 | 1 | 0 | 1 | 54 |

**Table S3.** Comparison of cell cluster classification of 19,344 hemocytes using 2,000 highly variable genes (eight clusters detected) and the 7,716 genes (10 clusters detected) that can be detected during library integration.

**Data S1.**
Log2 fold change in gene expression following selection for resistance to parasitic wasp and infection with parasitic wasp.

**Data S2.**

10x v2 library sequencing metrics generated from Cell Ranger count.

**Data S3.**

Enriched gene ontology categories and pathways for non-lamellocyte clusters. Enrichment analyses were performed on Flymine using the 2,000 highly variable genes as background.

**Data S4.**

Marker genes for lamellocyte differentiation. A random forest model was fit to identify the genes best able to predict the lamellocyte differentiation from the plasmatocyte progenitors. Markers are ranked by Gini impurity.

**Data S5.**

Enriched gene ontology categories and pathways for immature and mature lamellocyte clusters. Enrichment analysis were performed on Flymine using the 2,000 highly variable genes as background.

**Data S6.**

Genes detected in scRNA-seq dataset. 'All genes' corresponds to the list of genes detected in one or more scRNA-seq libraries. The top 2,000 highly variable genes in the integrated data sets were also identified.

**Data S7.**

Cluster markers identified from round one of clustering. The MAST test statistic was used to identify significantly upregulated markers in each cluster comparing the cluster of interest with cells in all other clusters. Pct.1 = percent of cells expressing genes in cluster of interest. Pct.2 = percent of cells expressing genes in all other clusters. avg_logFC = $\log_e$ fold change in expression in cluster of interest compared to all other clusters.

**Data S8.**

Cluster markers identified from round two of clustering. The MAST test statistic was used to identify significantly upregulated markers in each cluster comparing the cluster of interest with cells in all other clusters. Pct.1 = percent of cells expressing genes in cluster of interest. Pct.2 =

percent of cells expressing genes in all other clusters. avg_logFC = $\log_e$ fold change in expression in cluster of interest compared to all other clusters.

**Data S9.**

Cluster markers identified from round one of subclustering, using only lamellocytes and their plasmatocyte progenitor cells. The MAST test statistic was used to identify significantly upregulated markers in each cluster comparing the cluster of interest with cells in all other clusters. Pct.1 = percent of cells expressing genes in cluster of interest. Pct.2 = percent of cells expressing genes in all other clusters. avg_logFC = $\log_e$ fold change in expression in cluster of interest compared to all other clusters.

**Data S10.**

Marker genes for non-lamellocyte clusters. The MAST test statistic was used to identify significantly upregulated markers in each cluster. Significantly upregulated markers were identified by pairwise comparison to PLASM1. PLASM1 markers were identified by comparison with all plasmatocyte clusters (PLASM2, MET and AMP).
